# Optimal feedback solutions recapitulate key features of motor cortical population dynamics

**DOI:** 10.64898/2025.12.12.694046

**Authors:** Muhammad Noman Almani, Shreya Saxena

**Affiliations:** Center for Neurocomputation and Machine Intelligence, Wu Tsai Institute, Yale University, New Haven, CT, USA; Department of Electrical and Computer Engineering, Yale University, New Haven, CT, USA; Department of Biomedical Engineering, Yale University, New Haven, CT, USA

## Abstract

Neural populations display complex response patterns with marked transitions between distinct underlying computational strategies on very short timescales during motor tasks. Such complex-yet-structured dynamical strategies may reflect computational needs of neural systems, shaped by optimal feedback and autonomous mechanisms, in addition to biological constraints. Are there overarching computational principles that govern complex dynamical strategies exhibited by the neural population response? Here, we explore the hypothesis that computational strategies underlying neural population response represent optimal feedback solutions to the control of musculoskeletal dynamics through space for a goal. To validate this hypothesis, we develop a procedure called neural optimization using dynamical systems (NODS) learning to modify synaptic strengths within a recurrent network for locally-optimal feedback control of anatomically accurate musculoskeletal models during complex sensorimotor tasks. NODS learning works even when the objective function to be minimized is highly non-linear or the muscle model is very complex. The dynamical strategies underlying the neural network response constructed using NODS learning recapitulate key features of recorded population response. Importantly, optimal feedback solutions using NODS learning suggest that feedback mechanisms are essential for neural populations to flexibly transit between complex-yet-structured strategies. We further show that this framework provides theoretical foundations for why the solutions obtained using deep reinforcement learning algorithms extensively used to model sensorimotor tasks may explain the dynamical strategies underlying recorded population response. In summary, we develop novel methods and approaches suggesting that neural dynamics may be more strongly modulated by optimal feedback mechanisms, in addition to autonomous mechanisms, than previously appreciated.

## 1 Introduction

During voluntary limb movements and other motor tasks, the population responses in neural circuits exhibit complex-yet-structured activity patterns over time [1–7]. In motor cortex (MC), population trajectories evolve through distinct dynamical regimes that appear to serve different computational roles during preparation and movement [8–13]. A central hypothesis is that these strategies represent the computational demands of transforming sensory feedback into muscle excitations required to achieve a desired motor goal. This is plausible: there exist rich cortical pathways to MC from upstream brain regions that convey processed sensory feedback including proprioception [14–21]. The computational demands are further shaped by the biomechanics of the body [22–24], by circuit mechanisms given the existence of strong recurrent connectivity in MC [25, 26], as well as by evolutionary and biological constraints such as minimization of neural energy or muscle effort [27–36]. A key open question is whether there exist overarching computational principles that govern these sensorimotor transformations, and can explain the complex-yet-structured dynamical strategies observed in neural population responses.

Optimal feedback control (OFC) has provided a powerful normative framework for constructing behavioral-level models of biological movement [36]. It provides a principled way to obtain kinematic trajectories by positing that biological movements arise from solutions that optimize task performance, subject to biomechanical and energetic constraints [33, 37, 38]. In practice, however, OFC models of movement often employ simplified or linearized musculoskeletal dynamics, which yields tractable optimization, but sacrifices realism in the sensory feedback - particularly proprioception - that would impinge on MC [39, 40]. More recent optimal control methods can handle highly non-linear musculoskeletal models by focusing on the development of iterative procedures for locally-optimal control [41–43]. However, such approaches typically implement the controller as an abstract feedback law rather than as a realistic neural network. As a result, these approaches are well suited for explaining behavioral-level phenomena, but cannot directly account for the neural-level dynamical structure measured in MC.

In parallel, the machine learning community has developed powerful training algorithms for artificial neural networks to perform complex motor tasks. Feedforward architectures can generate sophisticated movements [44], and recurrent neural networks (RNNs) have been extensively used to model cortex-like dynamics with extensive recurrent connectivity [45, 46]. However, these approaches face fundamental challenges in their applicability to realistic sensorimotor control. Training RNN controllers for anatomically accurate musculoskeletal models involves highly non-linear dynamics and non-convex objectives, with no guarantees of converging to locally optimal solutions. Moreover, it is often difficult to interpret the resulting controllers through the lens of optimality principles, limiting their utility for testing normative theories of motor control.

Here, we bridge these perspectives by embedding a recurrent controller directly into the sensorimotor loop and training it using locally optimal feedback control. We introduce neural optimization using dynamical systems (NODS), which adapts iterative OFC algorithms [43] to construct time-varying linear dynamical systems that transform sensory feedback from anatomically accurate musculoskeletal models into muscle excitations required for delayed center-out reaches. The time-varying synaptic weights allow the linear models to implement effectively non-linear transformations of continuous sensory input while retaining much-needed interpretability. This framework preserves the normative appeal of OFC, operates directly on realistic musculoskeletal dynamics, and yields explicit predictions for the structure of neural population activity in MC during preparatory and movement epochs.

Many neural populations flexibly transition between distinct yet linked computational strategies across task epochs [47–51]. A canonical example is during a delayed reach, where recent work has shown that these computations occupy orthogonal-but-linked subspaces: preparatory and movement-epoch activity lie in nearly orthogonal subspaces that are nonetheless lawfully related [52–54] (Fig. 1A). Here, we ask whether such orthogonal-but-linked strategies naturally arise as optimal solutions for controlling realistic musculoskeletal systems. Using the proposed NODS framework (Fig. 1B), we show that controllers trained to perform delayed reaches not only achieve high kinematic accuracy but, importantly, recover key features of MC population dynamics, including pronounced orthogonality between preparatory and movement subspaces. These features cannot be captured using well-known motor cortical models [52].

**Fig. 1.**
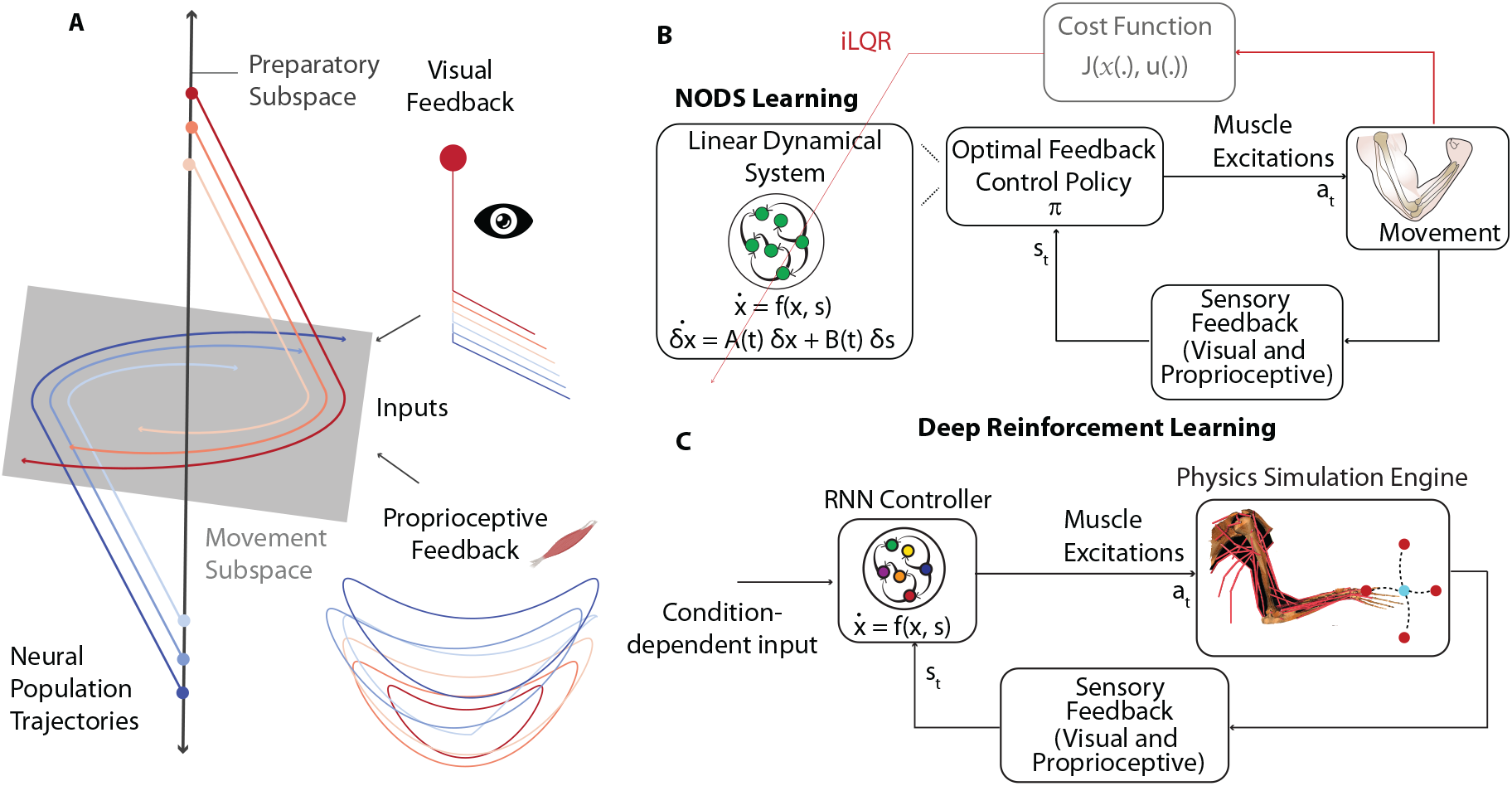
An illustration of orthogonality between the preparatory and movement population responses and the proposed frameworks for probing neural dynamics. **A**. An illustration of neural population trajectories in transition from preparation to movement during a delayed-reach task. During the preparatory period, the neural activity develops mainly within the single-dimensional preparatory subspace orthogonal to the two-dimensional movement subspace. On the movement onset, neural trajectories transition into the movement subspace and exhibit strong rotational dynamics. Each trace plots a neural population trajectory for one reach condition and color indicates the reach condition. In the proposed frameworks, we hypothesize that the motor cortex (MC) receives processed proprioceptive and visual feedback, in addition to condition-specific sparse inputs, as input presumably from upstream sensory processing regions. **B**. In NODS learning, optimal control policy *π* is formulated as a time-varying recurrent network with an internal state x_*t*_. Sensory feedback s_*t*_, consisting of proprioception and visual feedback, to the recurrent network is provided by an anatomically accurate musculoskeletal model. The recurrent network transforms the internal state x_*t*_ and sensory feedback s_*t*_ into the muscle excitation signals a_*t*_ through synapses of strength A(*t*) and B(*t*), which are modified using NODS learning. The anatomically accurate musculoskeletal model consisting of 6 muscles receives the generated muscle excitations and executes the corresponding movement. NODS learning modifies the synaptic strengths A(*t*) and B(*t*) to minimize a cumulative cost function *J* using iLQR algorithm. **C**. In DRL framework, the control policy or controller *π*, represents the transformation from sensory feedback s_*t*_ to muscle excitations a_*t*_ and is formulated using an RNN. The anatomically accurate musculoskeletal model of macaque arm consisting of 39 muscles is implemented in the MuJoCo physics simulation engine. Further training details using the cumulative reward or return *J* are given in Supplementary Fig. 1.

Finally, we connect this optimal control perspective to deep reinforcement learning (DRL), which has become a standard tool for training controllers for complex, high-dimensional musculoskeletal models. DRL policies are rooted in optimal control ideas, but are typically model-free and optimized for reward rather than explicit cost functions [55, 56]. We demonstrate that controllers trained with DRL on a realistic macaque arm model (Fig. 1C) exhibit similar dynamical motifs and subspace organization as the NODS-based controllers, while providing additional leverage to probe generalization and robustness to perturbations. By systematically manipulating proprioceptive and visual feedback, we show that rich sensory feedback, especially proprioception, is critical for online control and for the emergence of orthogonal-but-linked neural strategies. Together, these results suggest that optimal feedback mechanisms, interacting with realistic bodies and sensory information, may play a more prominent role in shaping cortical population dynamics than is captured by autonomous dynamical systems models.

## 2 Methods

### 2.1 Neural Optimization using Dynamical Systems (NODS)

The recurrent network that forms the basis of our optimal feedback control studies is a conventional time-varying linear dynamical system as defined by:

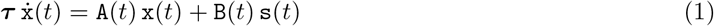

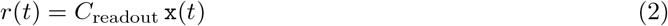

We define the muscle excitations or the instantaneous neural input as the readout of the recurrent network.

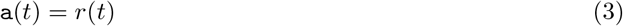

We use the terms ‘network’ and ‘controller’ interchangeably. Before specifying a task for the network, we first define its connectivity and output. The output of the individual neurons is characterized by firing rates. We define the network output as a weighted sum of firing rates or activities of its individual units. The recurrent network is a general purpose dynamical system that can be adapted for particular applications through connectivity and synaptic modification (Methods). The subsequent network connectivity and synaptic strengths are co-opted for any particular application using iterative optimal control algorithms (Methods). Although linear time-varying synaptic strengths and pre-defined readout weights are useful and meaningful ways to define the network for probing specific properties of neural dynamics as detailed below, it is a computational stand-in for complex sensorimotor circuitry. The mathematically elaborate nature of the iterative optimal control algorithms allows a flexible extension of the network architectures and nature of synaptic strengths to account for complex features of sensorimotor circuitry. Having defined the network connectivity and output, we now specify the task for the network to perform. The network receives specified sensory feedback, including proprioception, from the anatomically accurate musculoskeletal model (Methods) as a sensory input, s(*t*). The task for the network is then to generate muscle excitation signals, a(*t*), to perform specified movements. Here, it is important to note that we use anatomically accurate musculoskeletal models for generating accurate sensory feedback as an input for the network. In particular, proprioception requires that an accurate muscle model is implemented in musculoskeletal models. Such muscle models tend to have highly non-linear dynamics [43, 44]. This makes training of the network to generate movements using optimal feedback control algorithms challenging due to highly non-convex underlying objective function to be minimized. This challenge is usually resolved by instead incorporating a linear approximation of the musculoskeletal model. However, this is problematic because the generated sensory feedback is inaccurate, and further assumptions are explicitly imposed on the nature of feedback received by the biological MC to justify the results obtained and make contact with experimental data. The sensorimotor system can accomplish tasks in the presence of noise, delays, internal fluctuations and unpredictable environmental changes. Moreover, multiple lines of evidence suggest that motor cortical activity is optimized for musculoskeletal dynamics [22, 23]. This signifies the immense importance of the anatomical accuracy of musculoskeletal models and an elaborate feedback control scheme. Here, we overcome the challenge of incorporating anatomically accurate musculoskeletal models by using iterative OFC mechanisms to construct the controller for performing desired movements.

Consider a non-linear discrete-time dynamical system:

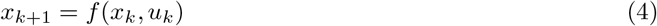

With

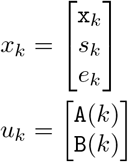

where, x is the controller’s activity or firing rates, *s* is the environmental or musculoskeletal model’s state and *e*_*k*_ = *g*(*x*_*k*_, *s*_*k*_) are the state variables corresponding to error signals purposely built for optimization procedures discussed below (Methods). Here, 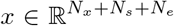 with *N*_*x*_, *N*_*s*_ and *N*_*e*_ corresponding to the number of controller’s units, sensory features, and error signal variables, respectively. 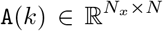 and 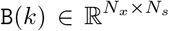 correspond to the recurrent and input synaptic strengths of the controller, respectively.

Sensory features, *s*, include proprioception, visual feedback and a go-cue signal. For notational clarity, the representation of go-cue in sensory features is suppressed. Proprioception consists of muscle lengths and velocities. Visual feedback consists of the difference between the *x* and *y* coordinates for the current hand position and the desired or target hand position. The error signals consist of the directional error between the current hand position and the target hand position:

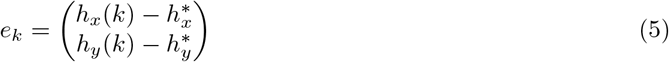

where *h*_*d*_ represents the current hand position and 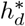 represents the target hand position. *d* ∈ {*x, y*} represents the *x* or *y* coordinate, respectively. The error signals are adjusted to construct controllers for different reaching conditions. A reaching target is specified by adjusting the corresponding 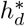 in the corresponding error signal.

Consider the instantaneous and final cost functions:

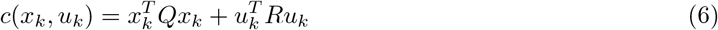

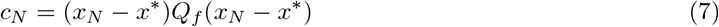

The cumulative quadratic cost function is:

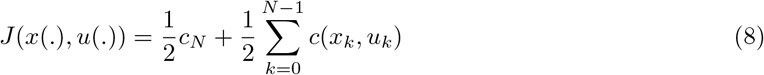

where *x*_*N*_ represents the final state and *x*^*∗*^ represents the desired target state.

Next, we follow Li and Todorov [43] to construct the optimal synaptic weights for the controller or inputs by linearizing the non-linear dynamical system (4) around the nominal trajectory. Each iteration begins with a nominal sequence *u*_*k*_ and a corresponding nominal state trajectory *x*_*k*_. An improved control sequence *u*_*k*_ is generated iteratively by linearizing the sensorimotor system dynamics around the nominal sequence *x*_*k*_, *u*_*k*_ and solving the modified linear quadratic regulator (LQR) problem. Let *δx*_*k*_, *δu*_*k*_ be the deviations from *x*_*k*_, *u*_*k*_. The linearized system dynamics are given by:

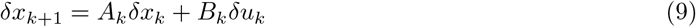

where *A*_*k*_ = *D*_*x*_*f* (*x*_*k*_, *u*_*k*_) and *B*_*k*_ = *D*_*u*_*f* (*x*_*k*_, *u*_*k*_). *D*_*x*_ is the Jacobian of *f* (.) w.r.t. *x* and *D*_*u*_ is the Jacobian of *f* (.) w.r.t *u*. The modified LQR problem has the cumulative cost function:

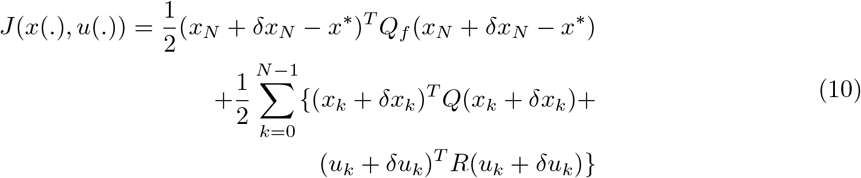

The following system of equations yield the optimal synaptic weights for the controller:

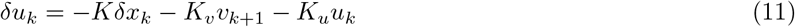

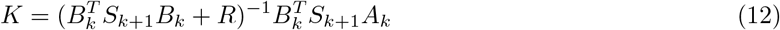

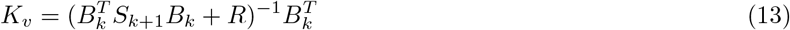

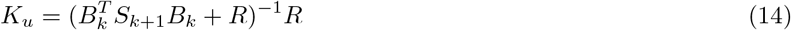

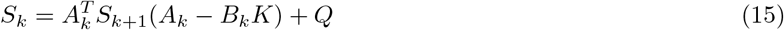

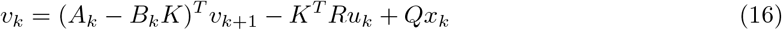

with the boundary conditions;

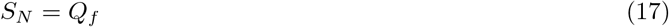

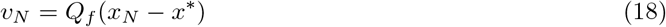

Because the algorithm modifies A and B for the locally optimal control of musculoskeletal systems based on the dynamics of both the controller and the musculoskeletal model, similar to the proposed function of neural controllers, we call it neural optimization using dynamical systems (NODS) learning. For notational clarity, we decomposed the matrices *Q* and *Q*_*f*_ to consist of the following block diagonal matrices:

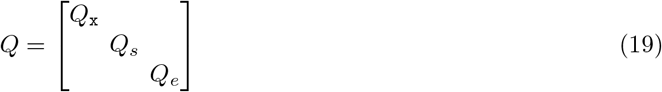

Similarly,

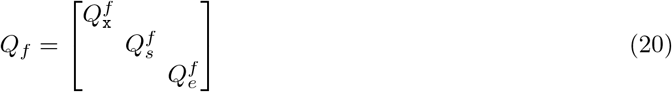

where *Q*_x_ and 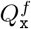 are the cost-weighting matrices corresponding to the network dynamics, *Q*_*s*_ and 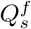 are the cost-weighting matrices corresponding to the sensory feedback (environmental and musculoskeletal dynamics), and *Q*_*e*_ and 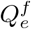 are the cost-weighting matrices related to the error-signal dynamics.

### 2.2 Connection with DRL

Recently, deep reinforcement learning (DRL) algorithms have become popular to train controllers to perform diverse movements using anatomically accurate musculokskeletal models. In the DRL framework, the sensorimotor loop formulation also consists of a controller interacting with and receiving sensory feedback from the environment (Fig. 1C). DRL shares rich connections with the optimal control framework. The theoretical foundations of DRL are, in fact, heavily based on the optimal control framework [57]. Analogous to the cost functions considered above, DRL is based on reward signals, *r*(*x*_*t*_, *u*_*t*_). Analogous to minimizing the cumulative cost function in OFC, DRL is based on maximizing the cumulative reward, *J*(*x*(.), *u*(.)), also known as the return [55].

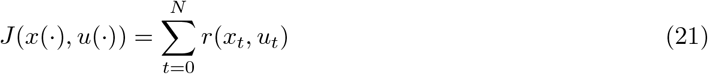

Oftentimes, a discounted return is considered for optimization purposes:

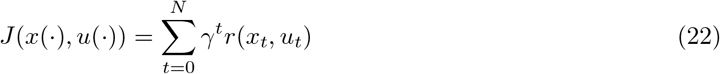

with the discounting factor, *γ* ∈ [0, 1]. The goal is to learn a policy or controller, *π*, that maximizes the return. Here, we consider a very specific formulation of the cumulative reward used in the soft actor critic (SAC) algorithm:

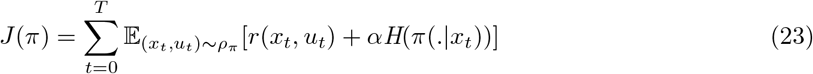

where, the temperature coefficient *α* determines the importance of the entropy term *H*(.) relative to the reward, and controls the stochasticity of the resulting policy. In OFC, the optimal policy is obtained analytically based on the underlying dynamics of the system as determined by (4). SAC is a model-free algorithm and does not depend on the system dynamics. The trained controller is obtained based on the exploration using a stochastic policy. In contrast to NODS learning, the policy obtained using DRL can generalize to conditions unseen during training [58]. In the next sections, we use the robustness to perturbation and generalization ability of DRL to generate key insights about the dynamical strategies underlying the neural population responses.

### 2.3 Implementation of the delayed-reach task

We followed the paradigm used in prior experimental studies [6, 13, 52] to construct the key events for the center-out delayed-reach task (Fig. 2). Relatively straight or slightly curved reaches were made between a central touch point and a radial outer target. Each simulated trial began with a baseline period. During the baseline period, the musculoskeletal model was initialized to begin with a random arm pose. We then used the NODS procedure detailed above to construct a controller to move the hand to a central fixation point. The error signals as well as the corresponding state and control cost-weighting matrices for the baseline period are given in Methods. During the baseline period, sensory feedback from the environment s(*t*) contained visual information of the central fixation location (besides the proprioceptive feedback). The baseline period was followed by the preparatory period. During the preparatory period, visual feedback of the target location was provided. Using the NODS scheme, a controller was constructed to stabilize the hand at the central touch point while preparing the reach to the indicated outer target (Methods). The preparatory period lasted for 375ms. The go-cue was set to 0 during this epoch. During the movement period, a go-cue (= 1) instructed the reach to the outer target. Visual feedback representing the target location was provided during the movement period. The movement period lasted for 200ms. Details of the error signals along with the state and control cost-weighting matrices used during the different epochs are given in Methods.

**Fig. 2.**
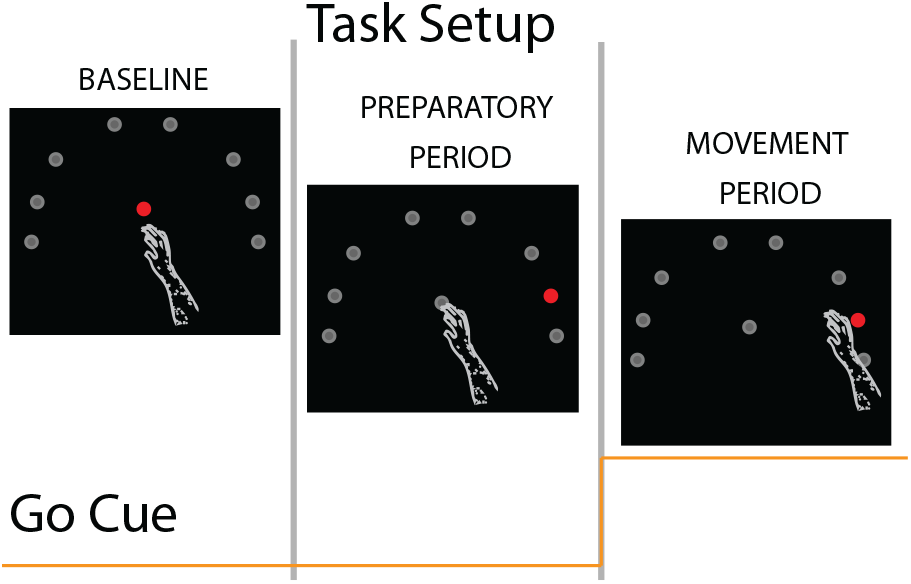
Events in the delayed-reach task implemented using the proposed frameworks. Controllers were constructed to perform delayed reaches to one of the eight possible target locations radially aligned roughly 10cm from the central fixation location. During the baseline period, sensory feedback, in addition to proprioception, contained visual information about the central fixation location (red circle). An additional go-cue signal in the sensory feedback was not active (=0). During the preparatory period, sensory feedback additionally contained visual information about the target location (red circle). Grey circles represent possible targets not represented in the sensory feedback. The go-cue signal remained inactive (=0) during the preparatory period. During the movement period, sensory feedback contained visual information about the reach target location (red). The go-cue signal remained active (=1) during the movement period.

## 3 Results

### 3.1 NODS controllers achieve high kinematic accuracy during reaches with a realistic musculoskeletal model

We simulated a center-out delayed-reach task for eight targets using the proposed NODS scheme. The controller acted on a two-joint six-muscle arm model (Methods) with the following muscles: shoulder flexor (SF), shoulder extensor (SX), elbow flexor (EF), elbow extensor (EX), biarticular flexor (BF), biarticular extensor (BX). Muscle parameters were based on anatomical measurements from macaque monkeys [59, 60].

The resulting movements reproduced a center-out delayed-reach task as shown in Figs. 3A-C. The targets were arranged radially at a 10 cm distance from a central fixation point. The simulated hand trajectories were smooth and nearly straight, with modest curvature that varied systematically with target location.

**Fig. 3.**
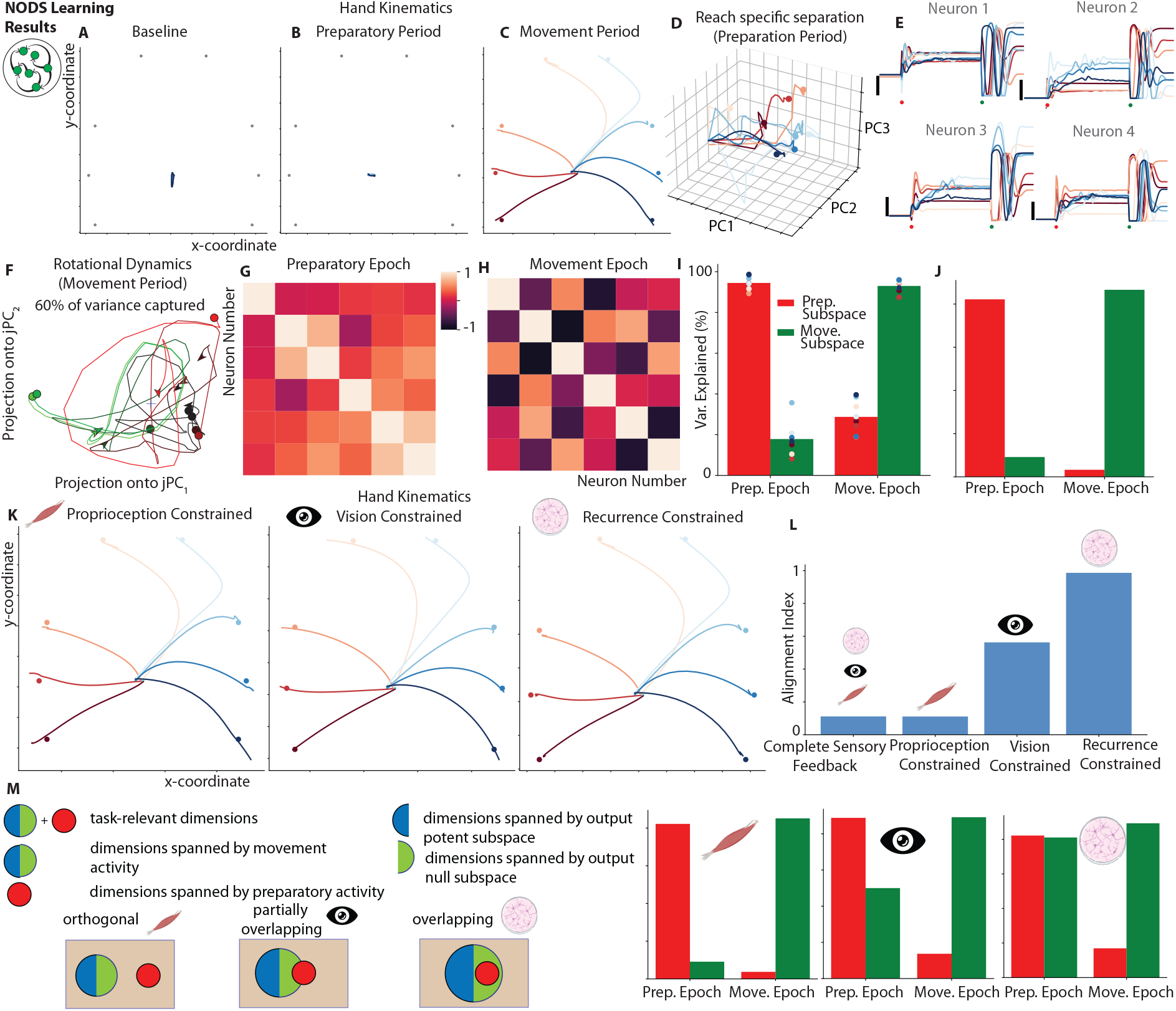
Controllers constructed with NODS learning suggest that feedback mechanisms are essential for constructed controllers to adopt the orthogonal-but-linked strategy. **A**. Hand trajectories (in the kinematic space) for the network constructed using NODS learning during the baseline period. **B** and **C**. Same as **A** but for the preparatory and movement periods, respectively. Trace color indicates the reach direction. **D**. Network population trajectories in the top three principal components (PCs) during the preparatory period. Circles indicate the end of the preparatory period. **E**. Responses of four example network neurons. Each trace is a network neuron’s firing rate during a reach in one of the eight directions. Trace color indicates reach direction (as indicated in **C**). Red dots represent the target onset time. Green dots represent the movement onset time. Black vertical bars represent a firing rate of 0.25 Arbitrary Units (AU). **F**. jPCA projections for the network firing rates data constructed using NODS learning during the movement period. Each trace shows the evolution of network state during the movement period for one of the eight directions. Circles denote the start of the movement period. **G**. Correlation matrix for the network neurons during the preparatory-epoch. **H**. Correlation matrix for the network neurons during the movement-epoch. The ordering of the network neurons is the same as for the preparatory-epoch correlation matrix. **I**. Percentage of variance captured by the preparatory (red) and movement (green) subspaces averaged across the eight reach conditions (colored circles). The left pair of bars corresponds to the variance captured during the preparatory-epoch. The right pair of bars corresponds to the variance captured during the movement-epoch. **J**. Same as **I** but for the cumulative neural response across eight reach directions. To calculate the cumulative neural response, we first concatenated the network’s response for each reach direction in the time dimension before computing the preparatory and movement subspaces. **K**. Movement period hand trajectories in the kinematic space for the proprioception, vision, and recurrence-constrained controllers. **L**. Alignment index for the unconstrained, proprioception, vision and recurrence-constrained controllers. **M**. Venn diagrams for the possible neural population structures (left-panel) and cumulative variance captured by preparatory and movement subspaces for the constrained controllers (right-panel).

### 3.2 NODS population dynamics recapitulate key motor cortical features

The NODS controller reproduced several features of MC population activity during delayed reaches. The preparatory activity developed shortly after the appearance of the target, during which population trajectories showed reach specific separation in the top principal components (Fig. 3D). Network activity transitioned from movement preparation to execution shortly after the go-cue instructed the reach. Single units exhibited temporally complex responses, and most were active during both preparatory and movement epochs. In the examples shown in Fig. 3E, the direction eliciting the highest firing rate during preparation rarely exhibited the highest firing rate during movement. During movement, population activity showed robust rotational dynamics as revealed by jPCA (Fig. 3F), consistent with empirical findings in MC [1].

To assess how structure reorganizes across epochs, we separately computed pairwise correlations for preparatory and movement epochs (Figs. 3G and H). The correlation structure changed markedly between the epochs for most units, other than those explicitly coupled in the model (Methods). Units that were strongly correlated during preparation did not necessarily maintain those relationships during movement, indicating a substantial reorganization of population-level structure and suggesting that activity perform different computations in the two epochs. This is difficult to reconcile with a simple view in which neurons with correlated responses are assumed to be functionally coupled in a way that generalizes across epochs. However, that view is rooted in models where MC dynamics are dominated by recurrent mechanisms. In feedback-driven framework, effective connectivity can be strongly shaped by input. The reorganization of correlations we observe therefore supports a picture in which feedback plays a central role in shaping population dynamics across task epochs.

### 3.3 NODS controllers implement orthogonal-but-linked preparatory and movement subspaces

The above observations suggest that preparatory and movement activity occupy distinct subspaces. Following [52], we considered three possible relationships between these subspaces. First, preparatory dimensions could be fully contained within movement dimensions that are output-null, such that preparation is gated by avoiding output-potent dimensions. Second, preparatory dimensions could partially overlap with the output-null movement dimensions. Third, preparatory and movement subspaces could be nearly orthogonal, sharing little or no overlap. We used principal components analysis (PCA) to distinguish between these possibilities (Figs. 3I and J). We performed PCA separately on the preparatory and movement-epoch network responses to identify preparatory PCs (prep-PCs) and movement PCs (move-PCs). Prep-PCs captured a large fraction of preparatory-epoch variance and move-PCs captured a large fraction of movement-epoch variance. However, prep-PCs captured very little movement-epoch variance, and move-PCs captured very little preparatory-epoch variance, indicating that the two subspaces are close to orthogonal.

We quantified the orthogonality with the alignment index defined by [52]. For the NODS controller, the alignment index was close to 0, indicating a very high degree of orthogonality between preparatory and movement subspaces. This level of orthogonality matches that observed in neural data and exceeds what has been reported for many existing recurrent models, including those based on a dynamical systems perspective [45, 52]. Thus, an optimal feedback controller operating in a realistic sensorimotor loop naturally adopts an orthogonal-but-linked strategy.

### 3.4 Rich sensory feedback is critical for orthogonal-but-linked strategies

Most existing dynamical systems models treat MC as an autonomous system driven by sparse, condition-dependent inputs rather than by rich, processed sensory feedback. Sparse inputs carry limited dynamical information, forcing the model to rely heavily on recurrent connectivity to produce complex activity patterns. In contrast, sensory feedback (especially proprioception) contains rich dynamical content during movement. This raises a key question: how important is processed sensory feedback for producing the orthogonality observed between preparatory and movement subspaces?

To address this, we constructed three types of constrained controllers using NODS (Methods). In proprioception-constrained controllers, only proprioceptive feedback was available. In vision-constrained controllers, only visual feedback was available. In recurrence-constrained controllers, the controller was forced to depend heavily on recurrent connections to generate motor output. All three types achieved high kinematic accuracy on the delayed-reach task (Fig. 3K).

Their population dynamics, however, differed markedly. Controllers constrained to use only proprioception exhibited the highest degree of orthogonality (lowest alignment index), followed by controllers that relied only on visual feedback (Fig. 3L). Controllers forced to depend heavily on recurrence showed the least orthogonality between preparatory and movement subspaces. These results indicate that rich dynamical content in the input promotes the emergence of orthogonal-but-linked strategies. They also imply that functional connectivity, and therefore correlations in response patterns, is strongly modulated by the structure of incoming sensory signals.

Consistent with this view, vision-constrained controllers exhibited partially overlapping population structures, while recurrence-constrained controllers showed strongly overlapping preparatory and movement subspaces (Fig. 3M). All of these structures were still compatible with preparatory activity avoiding output-potent dimensions, thereby preventing premature movement.

### 3.5 DRL-trained controllers achieve accurate reaches with a high-dimensional arm model

The NODS results highlight the power of optimal feedback mechanisms for explaining population-level strategies in the sensorimotor system. We next examined whether similar strategies emerge in controllers trained via DRL with a more realistic arm model.

Following Almani et al. [58], we simulated the delayed-reach task using the muSim framework. Specifically, we used an RNN-based controller with 256 hidden units (Methods) and an anatomically accurate musculoskeletal model of the macaque upper arm with 39 muscles (Methods). Sensory feedback included proprioception and visual feedback, matching the inputs in the NODS framework.

We trained the controller using the SAC algorithm with appropriate constraints and regularizations [58]. The trained controller achieved high kinematic accuracy for the delayed-reach task: the hand reamined near the central touch location during the preparatory period and reliably reached the cued target during the movement period (Fig. 4A).

**Fig. 4.**
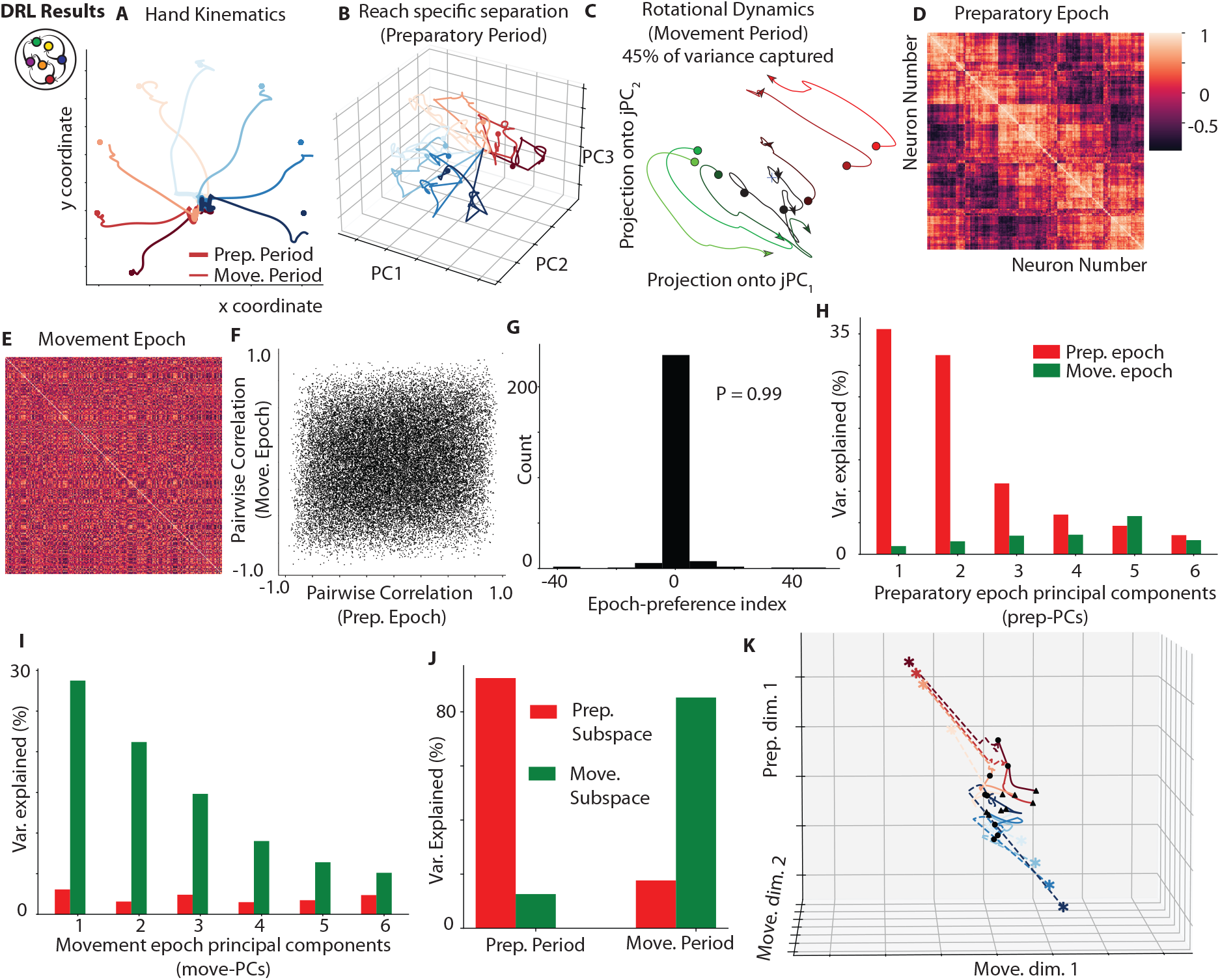
Controllers trained with DRL recover the orthogonal-but-linked strategy. **A**. Hand trajectories (in kinematic space) during the preparatory and movement periods for the controllers trained using DRL. **B**. Network population trajectories in the top three PCs during the preparatory period. Circles indicate the movement onset. **C**. jPCA projections for the network firing rates data trained using DRL during the movement period. Each trace shows the evolution of network state during the movement period for one of the eight directions. Circles denote the start of the movement period. **D**. Correlation matrix for the controller’s neurons during the preparatory-epoch. Each entry in the matrix denotes the strength to which the response pattern was similar for the two neurons during the preparatory-epoch. **E**. Correlation matrix for the controller’s neurons during the movement-epoch. The ordering of the network neurons is the same as for the preparatory-epoch correlation matrix. **F**. Correlation value for each network neuron pair during the movement-epoch is plotted against the correlation value for the same network neuron pair during the preparatory-epoch. **G**. Histogram corresponds to the epoch-preference index, which quantifies the strength of the controller’s activity during the preparatory-epoch compared with strength of the controller’s activity during the movement-epoch. The distribution is highly unimodal (Hartigan’s dip test; dip test statistic= 0.01; P= 0.99). **H**. Percentage of the preparatory-epoch data variance and movement-epoch data variance captured by the top six prep-PCs. **I**. Percentage of the preparatory-epoch data variance and movement-epoch data variance captured by the top six move-PCs. **J**. Percentage of the variance captured (for all eight reach conditions concatenated in the time dimension) by the preparatory (red) and movement (green) subspaces. The left pair of bars corresponds to the variance captured during the preparatory epoch. The right pair of bars corresponds to the variance captured during the movement epoch. **K**. Population trajectories from the trained controller emphasizing the transition from the preparatory to movement subspace. The trajectories are plotted in the top preparatory dimension and the top two movement dimensions. Black circles indicate movement onset.

#### 3.5.1 DRL population dynamics recover key features of MC activity

The DRL-trained controller also reproduced well-characterized features of MC population activity. During the preparatory period, trajectories in the top PCs showed target-specific separation (Fig. 4B). Flow fields during the delay period showed the presence of condition-dependent fixed-point (attractor) structure (Supplementary Fig. 2A). Single-unit responses also captured the key dynamical features as observed in the experimental data during the preparatory and movement epochs (Supplementary Fig. 2B). As in the NODS framework, jPCA revealed strong rotational dynamics during the movement epoch (Fig. 4C).

To examine how correlation structure reorganizes across epochs, we computed the correlation matrices for preparatory and movement activity (Figs. 4D and E, respectively). For the sake of visualization, units were ordered to highlight the correlation structure in the preparatory correlation matrix, and this ordering was reused for the movement matrix. Similarly to the OFC framework, the correlation structure changed significantly between the two epochs, with no systematic relationship between the correlations in the two epochs (Fig. 4F). Thus, the DRL controller also exhibits a pronounced reorganization of similarity structure across epochs.

We then asked whether this reorganization reflects different subpopulations specializing in each epoch. Following [52], we computed an epoch-preference index for each unit. The distribution was sharply peaked near zero (Fig. 4G), indicating that most units were strongly active in both epochs. As in the empirical data and the NODS framework, a largely overlapping set of units participates in both preparatory and movement computations, but their correlation structure and geometry reorganize across epochs.

#### 3.5.2 DRL controllers also implement orthogonal-but-linked subspaces

We next repeated the subspace analyses used for NODS. The top six prep-PCs captured a large fraction of preparatory-epoch variance but only a small fraction of movement-epoch variance (Figure 4H). Conversely, the top six move-PCs captured a large fraction of movement-epoch variance but little preparatory-epoch variance (Figure 4I). When preparatory and movement subspaces were computed on responses concatenated across reach directions, each subspace captured little variance from the other epoch (Figure 4J). As in the NODS framework, preparatory and movement-related computations occurred in nearly orthogonal subspaces.

Next, we examined a key question: how is the information flexibly transferred from the preparatory subspace to the orthogonal movement subspace? To visualize how information is transferred between these subspaces, we used the dimensionality reduction approach of Elsayed et al. (2016), which explicitly leverages subspace orthogonality. We identified a top preparatory dimension and top movement dimensions, and plotted trajectories in the space spanned by the top preparatory dimension and the top two movement dimensions (Figure 4K). Preparatory trajectories for different targets occupied distinct regions along the preparatory dimension. Around movement onset, trajectories left the preparatory dimension and entered the movement dimensions, while preserving the ordering of conditions. This is consistent with orthogonal-but-linked dynamics: preparatory activity sets initial conditions for movement-generation dynamics in an orthogonal subspace, while preserving target-specific information.

#### 3.5.3 Sensory ablation studies reveal proprioception is essential for the online control of movement

The emergence of orthogonal-but-linked strategies in both NODS and DRL frameworks points to a key role for sensory feedback. We therefore asked how sensory inputs contribute to the transfer of information between preparatory and movement subspaces.

The above findings raise another key question: what factors modulate the systematic transfer of information from one subspace to the other orthogonal subspace? One possibility, as indicated above, is that the processed sensory feedback may be a key factor in this transition by presumably modulating the functional connectivity between neurons across the two epochs. To examine this, we first performed sensory feedback ablation studies using the DRL framework (Fig. 5A). In the ablation studies, we selectively ablated proprioceptive or visual feedback to the controller 100ms before movement onset and observed the effect of this ablation on movement kinematics (Fig. 5B) and neural dynamics (Figs. 5C and D). We found that proprioceptive feedback ablation significantly disrupted movement period kinematics and the corresponding degree of orthogonality. Visual feedback ablation did not result in any significant disruption of movement kinematics or degree of orthogonality (Supplementary Fig. 3A). These findings suggest that proprioception is essential for the online control of movement regardless of orthogonality. This finding is consistent with the role of proprioception in movement generation as observed in the experimental studies [61–63]. However, visual feedback during the preparatory period may modulate relatively subtle or finer aspects of the evolution of neural population trajectories, rather than the degree of orthogonality. This echoes our NODS analyses in section 3.3, where proprioception strongly promoted high orthogonality and realistic neural strategies in both frameworks.

**Fig. 5.**
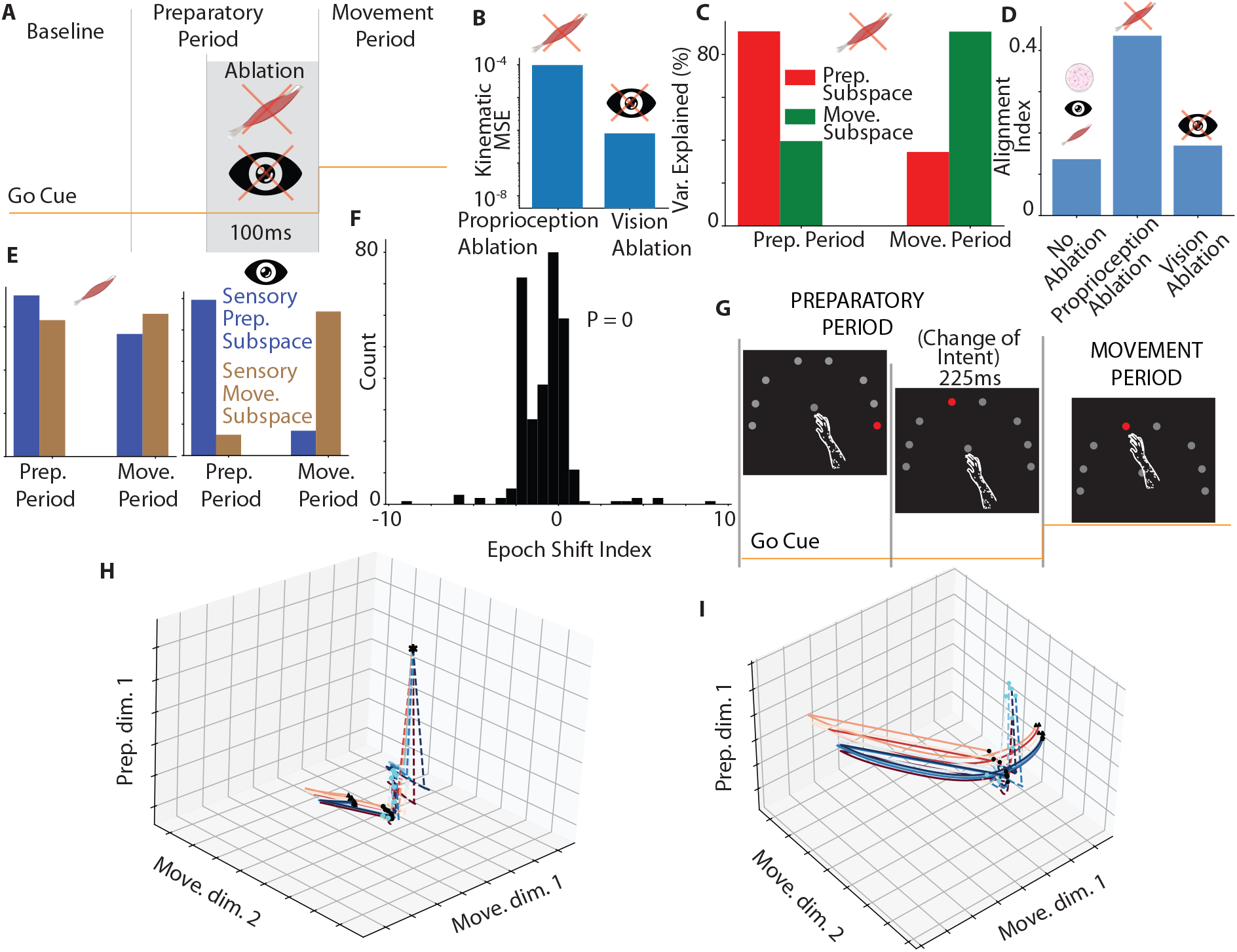
Sensory ablation and perturbation studies using DRL suggest that proprioception is essential for the online control of movement while visual feedback modulates the initial neural state for movement period dynamics. **A**. For the sensory ablation studies, either the proprioceptive or visual feedback to the trained controller was ablated 100ms prior to the movement onset. Icons indicate the type of sensory feedback that was ablated. **B**. Mean squared error (MSE) between the hand trajectories in the kinematic space obtained from controllers with different sensory feedback ablations and the corresponding hand trajectories obtained from the controller with no ablations averaged across all eight reach directions. **C**. Percentage of the variance captured by the preparatory (red) and movement (green) subspaces for proprioceptive feedback ablation. The left pair of bars corresponds to the variance captured during the preparatory-epoch. The right pair of bars corresponds to the variance captured during the movement-epoch. **D**. Alignment index for the unablated, proprioception-ablated, and vision-ablated controllers. **E**. Same as **C** but for the proprioceptive (left-panel) and visual (right-panel) feedback. **F**. Histogram corresponds to the epoch-shift index, which quantifies the strength of a sensory feedback feature being weighed by a controller’s neuron during the preparatory-epoch compared with strength of the same sensory feedback feature being weighed by the same controller’s neuron during the movement period. Positive values indicate that a controller’s unit is more selective towards a particular sensory feature during the movement period and negative values indicate that a controller’s unit is more selective towards a particular sensory feature during the preparatory period. The distribution is highly bimodal (Hartigan’s dip test; dip test statistic= 0.1; P= 0). **G**. For the sensory perturbation studies, the controller was presented with a target in one of the eight reach directions during the first half of the preparatory period. During the second half of the preparatory period, the target was perturbed and switched to the new location mandating the actual reach during the movement period. **H**. Population trajectories from the controller during sensory perturbation studies emphasizing the transition from the preparatory to movement subspace. The trajectories are plotted in the top-two movement dimensions and the top preparatory dimension. Cyan circles indicate attractors developed due to initially cued targets. Black circles denote the attractors developed due to the final cued target. Black triangles indicate 20ms after movement onset. **I**. The same space as in **H** rotated and zoomed to emphasize the phase-shift and separation between the movement-period population trajectories across different target switches. Black triangles indicate 50ms after movement onset.

To further disentangle the role of sensory feedback, we used a PCA procedure on sensory data similar to neural data to determine the orthogonality of the sensory feedback itself across the two epochs. The resulting subspaces after the PCA procedure on the sensory data during the preparatory and movement epochs are indicated as sensory preparatory and movement subspaces, respectively. Interestingly, we found that the proprioceptive feedback (as opposed to the visual feedback) was not orthogonal across the two epochs (Fig. 5E). Similarly, visual feedback had relatively distinct dynamical evolution of trajectories across the two epochs, as measured by dynamical similarity analysis (Supplementary Fig. 3B) (Methods). However, proprioception showed less trajectory tangling than visual feedback (Supplementary Fig. 3C). This suggests that dynamic content of proprioception, particularly across the two epochs, may provide an appropriate basis for the controller to adopt orthogonal-but-linked strategy.

#### 3.5.4 Single units reweight sensory features to switch between dynamical regimes

We then asked whether individual units flexibly reweight sensory features across epochs to support transitions between distinct dynamical regimes. To address this, we computed an epoch-shift index from the entries of the time-varying input weight matrix B(t) (Methods). An index near zero indicates similar weighting of a sensory feature in both epochs; positive values indicate stronger weighting during movement, and negative values indicate stronger weighting during preparation.

Many sensory features showed large positive or negative epoch-shift indices (Figure 5F), indicating that individual units substantially reoriented their sensory preferences across epochs. Thus, the dependence on sensory input is strongly time-varying, enabling complex activity patterns and smooth transitions between strategies. (Fig. 5F). In biological circuits, such flexibility may be implemented through nonlinear synaptic interactions between MC and upstream sensory processing regions [14], and may be modulated by go-cue signals transmitted to MC [10, 12].

#### 3.5.5 Sensory perturbations show visual feedback sets movement initial conditions

Finally, we examined how precise preparatory computations must be to correctly initiate movement generation dynamics. The optimal subspace hypothesis [8, 12] proposes that, during preparation, neural activity converges to a ‘prepare-and-hold’ region of state space that facilitates upcoming movement. This raises the question of how sensitive movement dynamics are to errors in the preparatory state, particularly in regimes where cortical mechanisms would be required to generate movement autonomously, without further sensory input.

We addressed this with change-of-intent perturbation experiments using the DRL controller (Fig. 5G). Here, during the first half of the delay period, visual feedback indicated one of the eight reach targets. During the second half of the delay period, the target switched to a different location, and the controller was required to reach the new target during movement. Despite the incorrect initial cue, the controller reliably executed accurate reaches to the final target (Supplementary Fig. 4).

Next, we examined changes in the neural population trajectories following incorrect motor preparation caused by the target switch. We found that during the first half of the preparatory period, population trajectories approached an attractor for the initially cued reach (Figs. 5H and I). However, the target switch induced a marked change in the evolution of network population trajectories. After the switch, the trajectories departed and converged to a new attractor corresponding to the final target location. The position of the new attractor depended on the initial target, indicating that the prepare-and-hold region is broader than often appreciated: a family of preparatory states can seed successful movements.

Movement-generation dynamics continued to unfold in a subspace orthogonal to the preparatory subspace, but with a clear phase shift between movement-period rotational dynamics across different target-switch conditions. This finding suggests that aspects of movement-generation dynamics depend on the particular preparatory state prior to the switch, possibly to correct residual errors in the initial neural state. The phase shift persisted for *∼* 110 ms into the movement period and was especially pronounced along the preparatory dimension, suggesting that movement-generating components remained relatively conserved. These observations are consistent with the idea that slightly different movement-generating dynamical systems, each relying on ongoing proprioceptive feedback and the final target, are engaged depending on the preparatory state, while visual feedback primarily shapes the initial neural states for movement-generation dynamics.

### 3.6 Discussion

We set out to ask whether the striking structure of motor cortical population activity during delayed reaches can be understood as a consequence of optimal feedback control operating on realistic bodies and sensory streams. Using NODS, a locally optimal feedback framework that learns time-varying linear recurrent controllers for an anatomically grounded musculoskeletal arm, we found that optimal solutions naturally reproduce key motifs reported in motor cortex: low-dimensional trajectories, strong rotational dynamics during movement, reorganization of pairwise correlations across epochs, and nearly orthogonal but lawfully linked preparatory and movement subspaces. Recurrent policies trained with DRL on the same embodied task converged on similar dynamical strategies, supporting the idea that these motifs are natural solutions to an embodied control problem rather than fragile properties of a particular model class.

#### Optimal feedback and the origin of orthogonal-but-linked dynamics

Empirical work suggests that preparatory activity occupies one subspace, movement-related dynamics unfold in another nearly orthogonal subspace, and there exists a mapping between them. Dynamical systems models can reproduce parts of this picture but often require finely tuned recurrence and still fall short of the strong orthogonality seen in data. In our framework, orthogonal-but-linked dynamics emerge directly from solving an online control problem under biomechanical and sensory constraints. Preparation organizes initial conditions in a way that is informative about the upcoming movement while avoiding output-potent dimensions; movement-related dynamics then unfold in a subspace optimized for stabilizing trajectories and correcting perturbations.

#### Sensory feedback, especially proprioception, shapes population structure

Our results highlight sensory feedback, not just recurrent circuitry, as a major determinant of cortical geometry. NODS controllers constrained to use only proprioception still achieved high kinematic accuracy and exhibited the strongest orthogonality between preparatory and movement subspaces, whereas recurrence-dominated controllers showed more overlapping subspaces and less clean rotational structure. DRL-based sensory ablations mirrored this: removing proprioceptive feedback strongly disrupted both kinematics and subspace geometry, while removing visual feedback had modest effects on these metrics. Analysis of the sensory signals themselves showed that proprioceptive trajectories are smoother and less tangled across epochs than visual feedback, providing a natural basis for separating ‘initialization’ and ‘execution’ computations. In this view, changes in pairwise correlations and subspace organization across epochs reflect changes in how rich sensory features are weighted over time, rather than a hand-off between distinct hard-wired subnetworks.

#### Convergence of NODS and DRL

Although NODS and DRL differ in objectives and training procedures, both arrived at similar dynamical strategies when given the same body and sensory inputs: accurate reaches, rotations during movement, overlapping sets of units active across epochs, and orthogonal-but-linked subspaces. In NODS, this flexibility appears as time-varying input weights that linearly reweight proprioceptive and visual features across time; in DRL, it appears as units that non-linearly transform proprioceptive and visual features across time. This convergence suggests that these population motifs are robust solutions to the embodied control problem. It also implies that task-trained networks can serve as testbeds for hypotheses about neural geometry, while OFC-inspired analyses help interpret how changes in cost, biomechanics, or sensory bandwidth should reshape subspaces and dynamics.

#### Limitations and future directions

Our models make several simplifying assumptions: NODS uses time-varying linear dynamics; sensory preprocessing is simplified; and the task involves relatively simple delayed reaches. Extending these frameworks to nonlinear OFC-inspired architectures, richer sensory transformations, and more complex behaviors (multi-step actions, posture, object interaction, social contexts) will test how general orthogonal-but-linked strategies are and how additional circuits (e.g., basal ganglia, cerebellum) interact with cortical controllers. A further challenge is to connect these optimal solutions to biologically plausible learning rules and developmental processes, explaining not only why such strategies are good, but how the brain might discover them.

In summary, by combining NODS and DRL in realistic musculoskeletal models, we show that orthogonal-but-linked subspaces, rotational dynamics, and epoch-dependent correlation structure can emerge from the demands of controlling a real body with rich sensory feedback. Proprioception plays a central role in enabling online control and shaping population geometry, while vision primarily sets initial conditions for movement-period dynamics. This offers a unified, embodied account of motor cortical activity and concrete predictions for how neural dynamics should change under manipulations of feedback, task structure, or biomechanics.

## 4 Extended Methods

### 4.1 Musculoskeletal model for OFC studies

We considered a two-joint arm model with shoulder and elbow joints. The joint torque *τ* ∈ ℝ^2^ is given as:

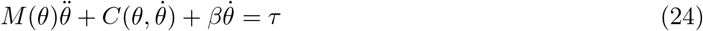

where, *θ* ∈ ℝ^2^ is the joint angles vector (*θ*_1_: shoulder angle ; *θ*_2_: elbow angle). 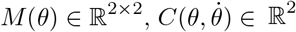 and *β* ∈ ℝ^2×2^ are the inertia matrix, centripetal and Coriolis forces vector and joint friction matrix, respectively.

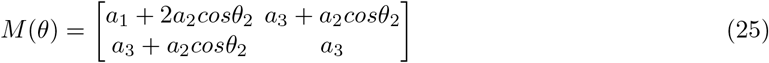

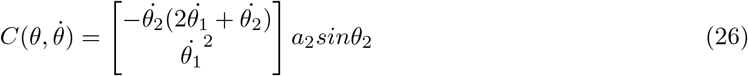

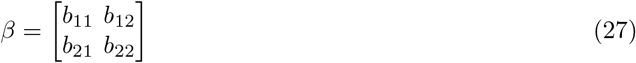

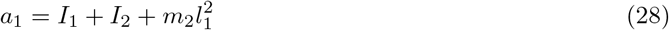

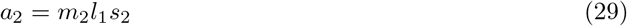

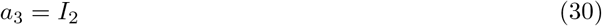

where *b*_11_ = *b*_22_ = 5 × 10^−3^, *b*_12_ = *b*_21_ = 2.5 × 10^−3^ represent joint friction. *m*_1_ = *m*_2_ = 0.3 (kg) represent the mass of the link *i* (*i* = 1, upper arm; *i* = 2 forearm). *l*_1_ = 0.15m and *l*_2_ = 0.21m represent the length of link *i. s*_*i*_ represent the distance from mass center for link *i*, with *s*_1_ = 0.07 and *s*_2_ = 0.12 (m). *I*_*i*_ is the moment of inertia with *I*_1_ = 5 × 10^−3^ and *I*_2_ = 9 × 10^−3^. We considered the following muscles: shoulder flexor (SF), shoulder extensor (SX), biarticulate flexor (BF), biarticulate extensor (BX), elbow flexor (EF), elbow extensor (EX). The moment arms matrix *A* ∈ R^2×6^ is given by:

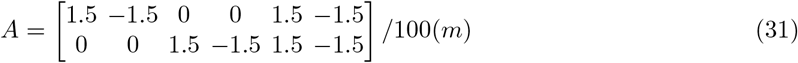

All the parameters were estimated from measurements of a macaque monkey [59, 60, 64]. The normalized muscle lengths were calculated as:

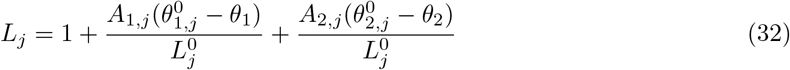

The muscle velocities are given as:

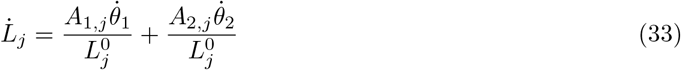

where subscripts index into the relevant matrix or vector. *θ*^0^ represent the optimal joint angle vector corresponding to the maximal joint torque generation. We considered the following *θ*^0^.

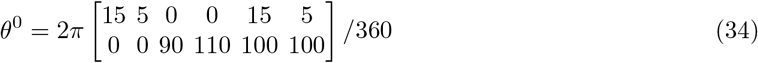

The optimal muscle lengths were given as:

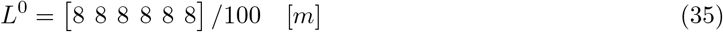

Joint torque *τ* is given as a product of moment arms *A* and the muscle tension vector *T* ∈ *ℝ*^6^:

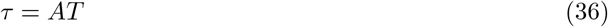

We modeled the torque due to gravity as done in previous works [65]:

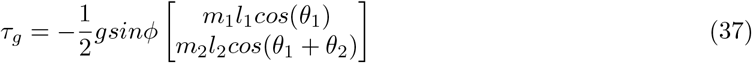

where, *ϕ* = 90° denotes the inclination angle of the workspace, and *g* = 9.8*ms*^−2^ is the acceleration due to gravity. The forward dynamics of the arm model are then given by:

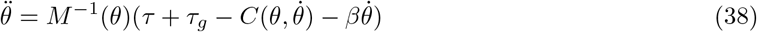

The jth muscle activation *a*_*j*_ is generated by passing the instantaneous neural input a_*j*_ through a filter that describes calcium dynamics. We modeled this using a first order nonlinear filter of the form:

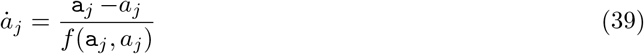

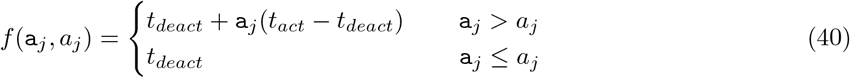

where, *t*_*act*_ = 0.05(*s*) and *t*_*deact*_ = 0.066(*s*) reflect faster input-dependent activation patterns than constant deactivation patterns.

The *jth* muscle tension *T*_*j*_ is calculated by scaling the corresponding unit-less tension 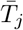 with the absolute muscle force (*F*_*a*_) of the physiological cross-sectional area (PCSA).

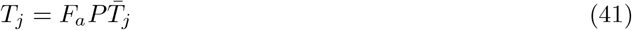

where *j* indexes into the relevant vector. *F*_*a*_ = 32(*N/cm*^2^) is the absolute muscle force and *P* = 10(*cm*^2^) is the PCSA assumed to be constant across the muscles. The unitless muscle tension is given as the sum of the contractile element (*F*_*CE*_(.)) and passive elastic element (*F*_*P E*_(.)).

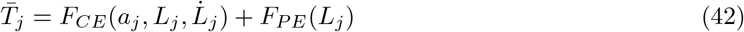

where,

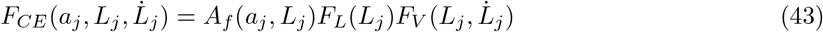

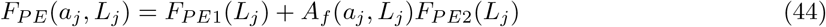

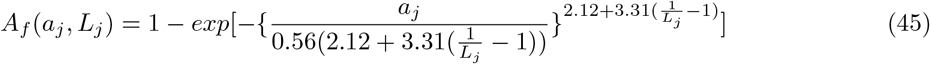

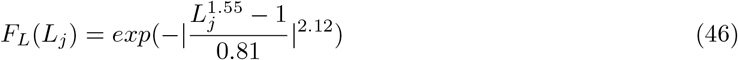

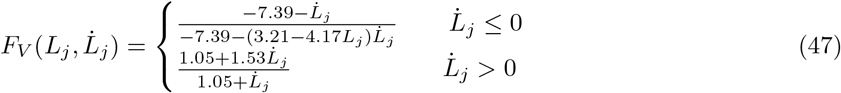

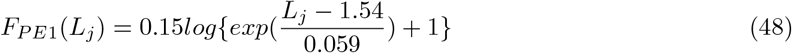

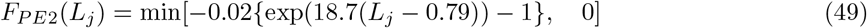

Finally, the transformation from the joint positions and velocities to hand positions and velocities is given as:

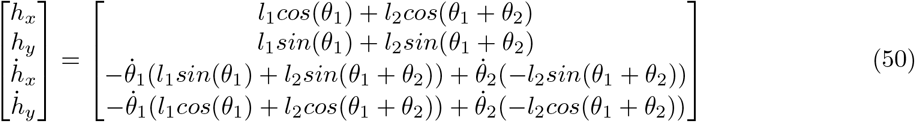

### 4.2 Network architecture for OFC studies

For the OFC studies, we modeled the RNN as a time-varying linear dynamical system:

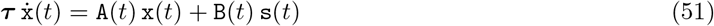

where, s is a vector consisting of proprioceptive (*L*, 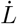) and visual feedback. Visual feedback was modeled as the distance between the hand coordinates (*h*_*x*_ and *h*_*y*_) and the target coordinates 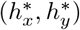. We formulated two error signals (*e*_1_ and *e*_2_) to model the visual feedback during the three key events in the delayed reach task. The instantaneous neural activations or muscle excitations a_*t*_ were given as the readout of the time-varying linear dynamical system.

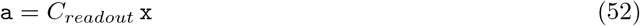

where, x is the vector of the network’s firing rates with dimensionality 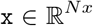. We used *N*_*x*_ = 22. We assumed that half of the network neurons directly modulated a particular muscle activation with the corresponding entries in the *C*_*readout*_ set to 1 for a mapping to a particular muscle. We assumed that the remaining half of the network neurons co-modulated all the muscle activations with the corresponding entries set to 1 in the *C*_*readout*_ matrix for all 6 muscles.

### 4.3 Optimal feedback control and formulation of error signals

After formulating the forward dynamics, we define the environmental or musculoskeletal state *s* corresponding to the state variables:

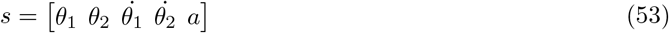

To model visual feedback during the delayed-reach task, we formulate the following error signals:

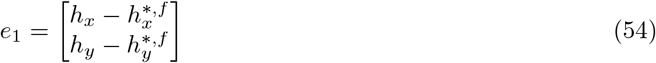

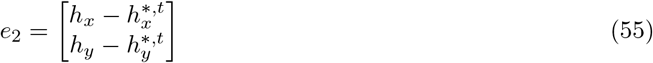

where, *h*^*∗,f*^ correspond to the coordinates of the central fixation point and *h*^*∗,t*^ correspond to the coordinates of the target location. Therefore, the system state was defined as:

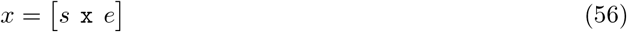

The cost-weighting matrix *Q*_*s*_ is decomposed into block-diagonal matrices for convenience.

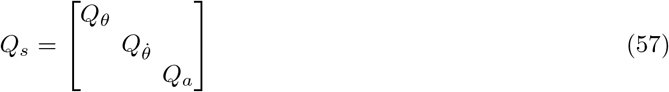

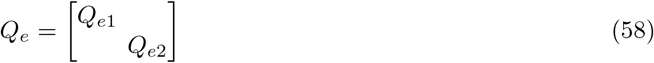

We follow a similar block diagonal decomposition for the final state-cost weighting matrices, 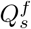 and 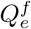.

The input cost-weighting matrix is also decomposed into block diagonal matrices as detailed below:

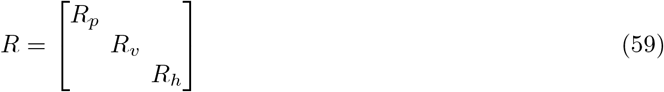

where 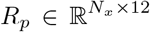 corresponds to the inputs or network weights associated with proprioceptive feedback, 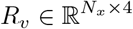 corresponding to the network weights associated visual feedback and 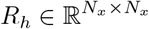 corresponding to the recurrent weights.

### 4.4 Cost-weighting matrices and error signals for key events in the delayed-reach task

To model key events, i.e. baseline, preparatory and movement periods, in the delayed reach task, we used specific cost weighting matrices corresponding to each event. During optimization using NODS learning, the final state from the baseline period was used to initialize the system state for the preparatory period. Similarly, the final state from the preparatory period was used to initialize the system state for the movement period. This is consistent with the Bellman’s principle of optimality which states any part of the optimal trajectory is also optimal. All the matrices were assumed to have proper dimensions and are not indicated below for notational clarity. We used 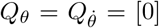 for all the key events. Similarly, we used *Q*_*a*_ = [*I*] × 10^−8^ and *Q*_x_ = [*I*] × 10^−4^ for all the three key events. For the baseline period, we used *Q*_*e*1_ = [*I*] × 15 × 10^3^, *Q*_*e*2_ = [0] and *R*_*p*_ = *R*_*v*_ = *R*_*h*_ = [*I*] × 10^1^. For the preparatory period, we used *Q*_*e*1_ = [*I*] × 9 × 10^8^, *Q*_*e*2_ = [*I*] × 3 × 10^6^ and *R*_*p*_ = *R*_*h*_ = *R*_*v*_ = [*I*] × 5 × 10^0^. For the movement period, we used *Q*_*e*1_ = [0], *Q*_*e*2_ = [*I*] × 6 × 10^3^ and *R*_*p*_ = 5 × 10^−2^ *R*_*v*_ = *R*_*h*_ = [*I*] × 5 × 10^0^.

### 4.5 Cost-weighting matrices for the constrained network models

For constructing the constrained network models, only the input cost-weighting matrices were adjusted. All the other cost-weighting matrices remained the same as detailed above. For the proprioception constrained models, we used *R*_*p*_ = [*I*] × 5 × 10^−2^ and *R*_*v*_ = *R*_*h*_ = [*I*] × 10^21^. For the vision constrained models, we used *R*_*v*_ = [*I*] × 5 × 10^−2^ and *R*_*p*_ = *R*_*h*_ = [*I*] × 10^21^. For the recurrence constrained models, we used *R*_*h*_ = [*I*] × 5 × 10^−2^ and *R*_*p*_ = *R*_*v*_ = [*I*] × 10^21^.

### 4.6 Musculoskeletal model for DRL studies

We use an anatomically accurate musculoskeletal model as developed in [58]. The model consists of 39 muscles. These muscles are based on the anatomical data obtained from cadaveric studies [66]. The musculoskeletal model consists of seven degrees of freedom (DoF) that include extension and flexion of the elbow, 3D rotation about the shoulder joint, supination and pronation of the lower forelimb, adduction/abduction and flexion/extension of the wrist. The five segments represented in the model are the hand, the torso, the radial side of the lower arm, the ulnar side of the lower arm and the upper arm. Further details on the model are given in [58].

### 4.7 RNN architecture for DRL studies

The policy network for the DRL studies consisted of three layers. The first layer was a feedforward layer followed by an RNN layer. The final feedforward layer represented the transformation to the muscle excitation signals. Neural regularizations on the policy network and additional hyperparameter details used during training are detailed in [58].

### 4.8 Epoch shift index

We calculated the strength of the tuning of the controller’s neuron *i* for a particular sensory feature *j* separately during the preparatory and movement-epochs. First, we calculated the mean weighting (across all reach conditions and times) of sensory feature *j* by the controller’s neuron *i* during the preparatory period using the entries of the matrix *B*(*t*) in (1), *S*_PREP_(*i, j*). Similarly, we calculated the mean weighting of the sensory feature *j* by the controller’s neuron *i* during the movement period, *S*_MOVE_(*i, j*). To account for the fact that preparatory and movement sensory weightings may have different average magnitudes for a given neuron *i*, we normalized the tuning of neuron *i* for the sensory feature *j* by the mean across all sensory features *j* for a given neuron *i* separately during the preparatory 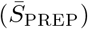 and movement-epochs 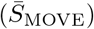. The epoch shift index *S*(*i, j*) of neural activity for a controller neuron *i* and sensory feature *j* is then:

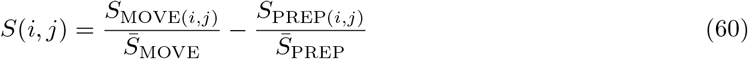

This index measures the preference for the tuning of a neuron *i* towards a sensory-feature *j* during preparatory versus movement epochs. If the resulting distribution of this index peaks at some value other than 0, it signifies that there exists a subpopulation with a particular preference for sensory features during the preparatory-epoch and a reoriented preference for sensory features during the movementepoch.

### 4.9 Dynamical Similarity Analysis

As the name suggests, dynamical similarity analysis (DSA) measures the dynamical similarity between the trajectories sampled from two dynamical systems. Consider two dynamical systems:

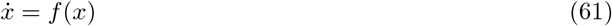

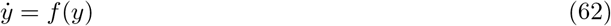

where *X* ∈ ℝ^*t*×*n*^ and *Y* ∈ ℝ^*t*×*n*^ represent trajectories sampled from the two dynamical systems, respectively, with *t* is the number of timepoints, and *n* is the number of state variables for the dynamical systems. Delay-embedded Hankel tensors with lag *p* are created *H*_*x*_ ∈ ℝ ^(*t*−*p*)×*n*^. Dynamic mode decomposition (DMD) matrices are created based on hankel alternative view of koopman (HAVOK). Procrustes analysis is then used to measure the alignment between the DMD matrices through a similarity transformation and report the final DSA score. A high DSA score indicates high similarity between the trajectories from the two dynamical systems. A low DSA score indicates otherwise.

### 4.10 Trajectory Tangling

Trajectory tangling was quantified using:

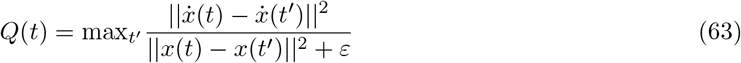

where *x*(*t*) is the system’s state at time 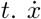 represents its temporal derivative and was calculated Using 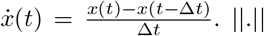. ||. || represents the Euclidean norm and *ε* was set to a very small value to prevent the division by 0. Further details on the hyperparameters used are given in [67].

### 4.11 Supplementary Figures

**Supplementary Figure 1.**
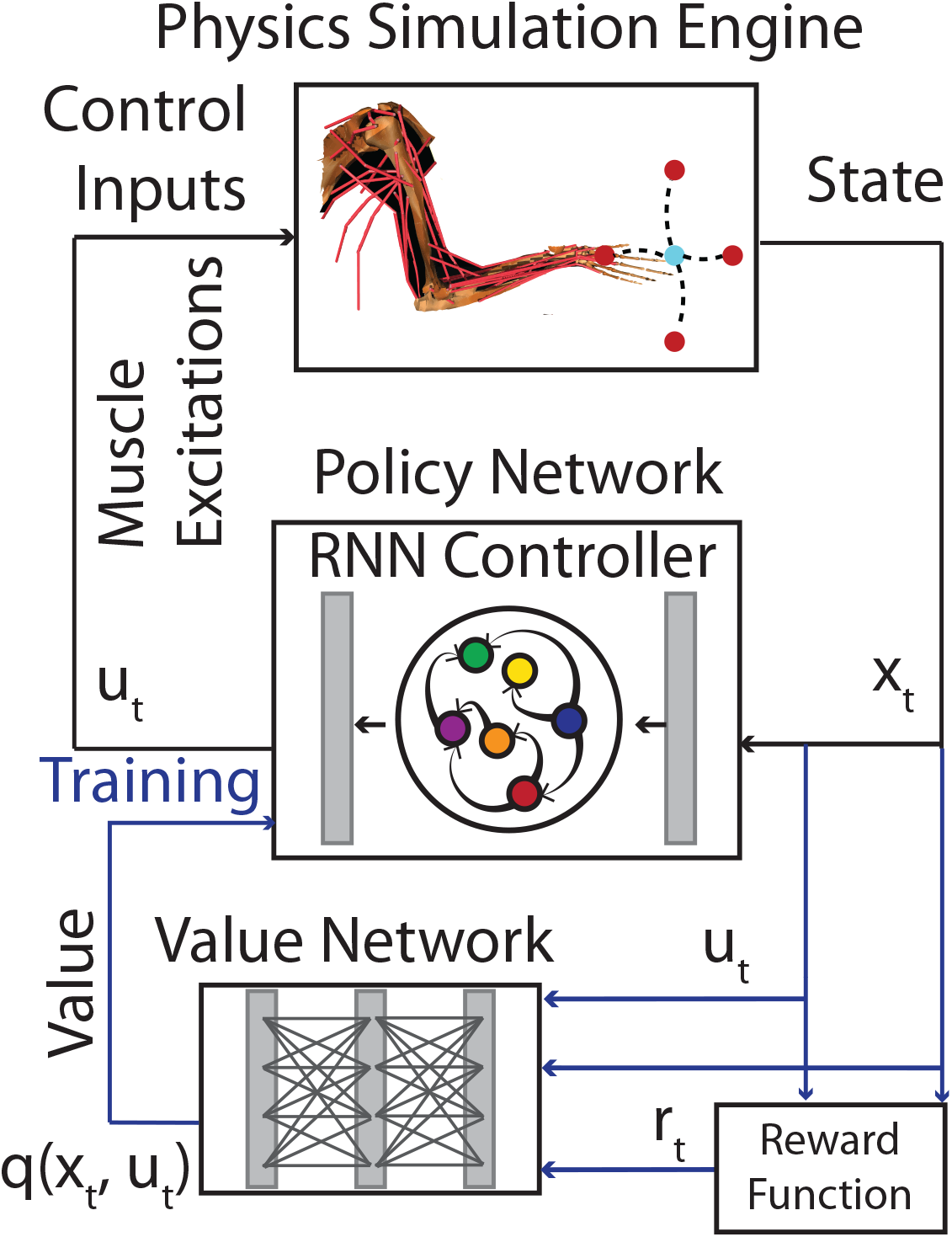
Training details for the DRL framework. In DRL, cumulative reward (analogous to cumulative cost in the NODS framework), also known as return *J*, is used to train the controller. Reward function is specified as a function of the state *x*_*t*_ and muscle excitations *u*_*t*_, *r*(*x*_*t*_, *u*_*t*_). During training, the cumulative reward function is parameterized using the value network. Value network represents the cumulative reward or return, *q*(*x*_*t*_, *u*_*t*_), for the corresponding state and action pair. Black connections are used during training and testing. Blue connections are used only during training.

**Supplementary Figure 2.**
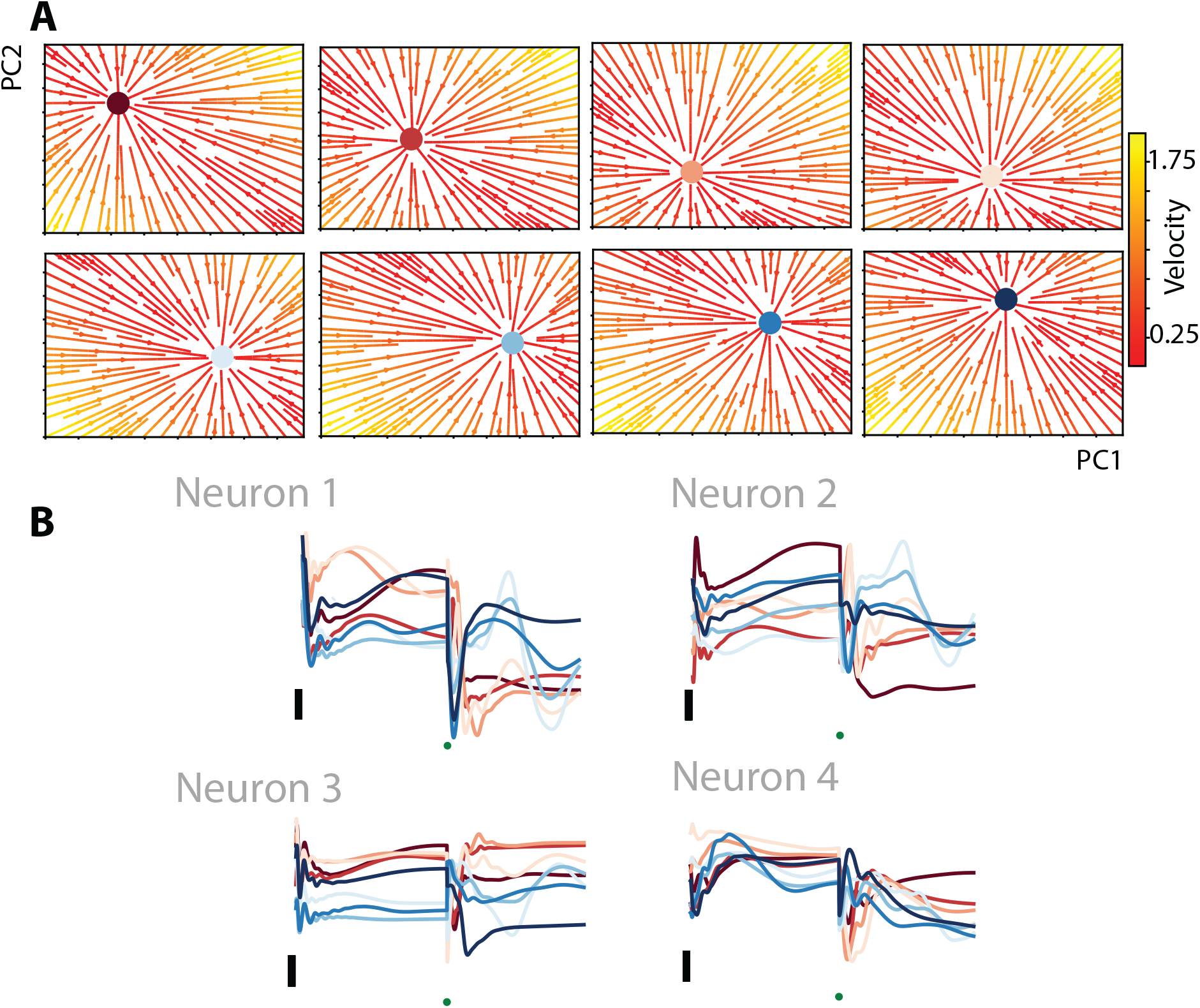
Flow-fields for sensory perturbation and single-neuron responses for DRL framework. **A**. Flow fields for the network response for the eight reach directions during the preparatory period in the top 2 PCs subspace. To construct the flow fields, the sensory feedback in the original high-dimensional sensory space was perturbed. The network trajectories converged to the reach target specific attractors (colored circles) after perturbation. **B**. Responses of four example network neurons. Each trace is a network neuron’s firing rate during a reach in one of the eight directions. Trace color indicates reach direction. Green dots represent the movement onset. Black vertical bars represent a firing rate of 0.025 Arbitrary Units (AU).

**Supplementary Figure 3.**
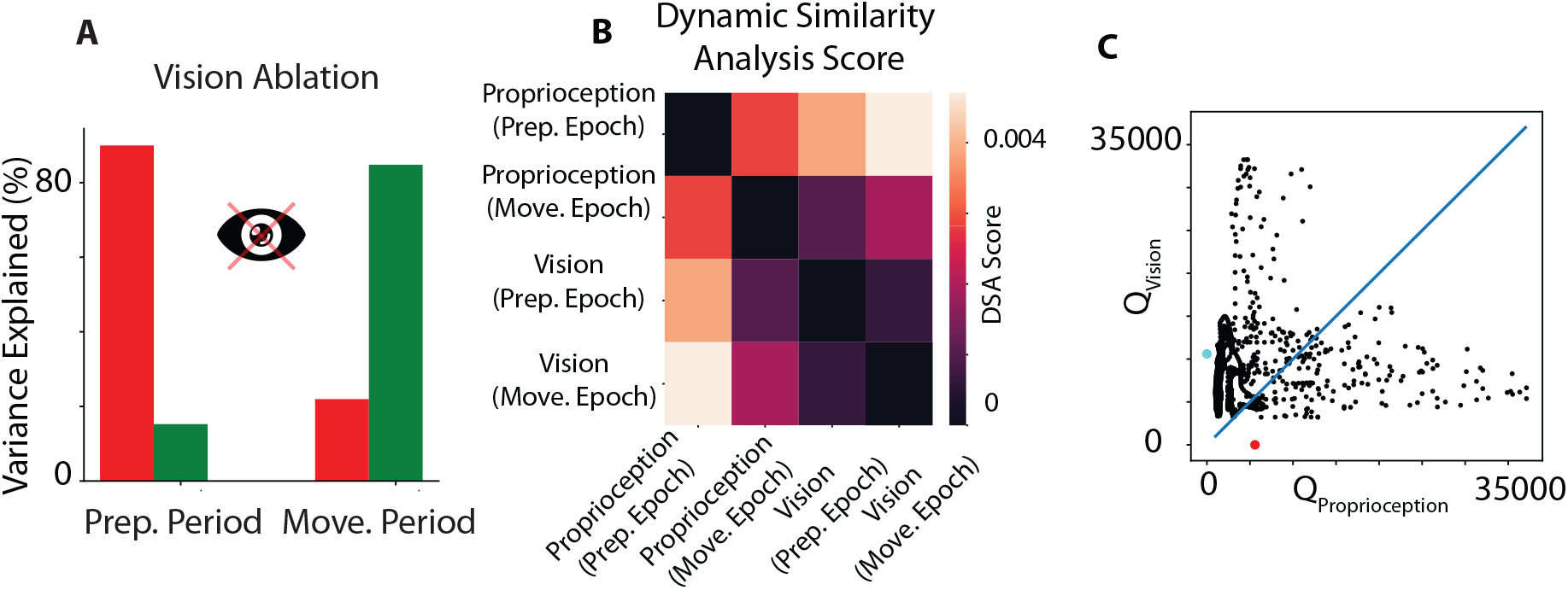
Orthogonality for visual feedback ablation and dynamical characteristics for sensory feedback. **A**. Percentage of the variance captured by the preparatory (red) and movement (green) subspaces for visual feedback ablation. The left pair of bars corresponds to the variance captured during the preparatory epoch. The right pair of bars corresponds to the variance captured during the movement epoch. **B**. Dynamical similarity analysis (DSA) score matrix for the proprioceptive and visual feedback during the preparatory and movement-epochs. **C**. Trajectorytangling values for the visual feedback (*Q*_Vision_) plotted against the proprioceptive feedback (*Q*_Proprioception_) during the movement-epoch. Each black dot represents the trajectory-tangling value for a particular timepoint. Cyan and red circles indicate 85th percentile of respective trajectory tangling values.

**Supplementary Figure 4.**
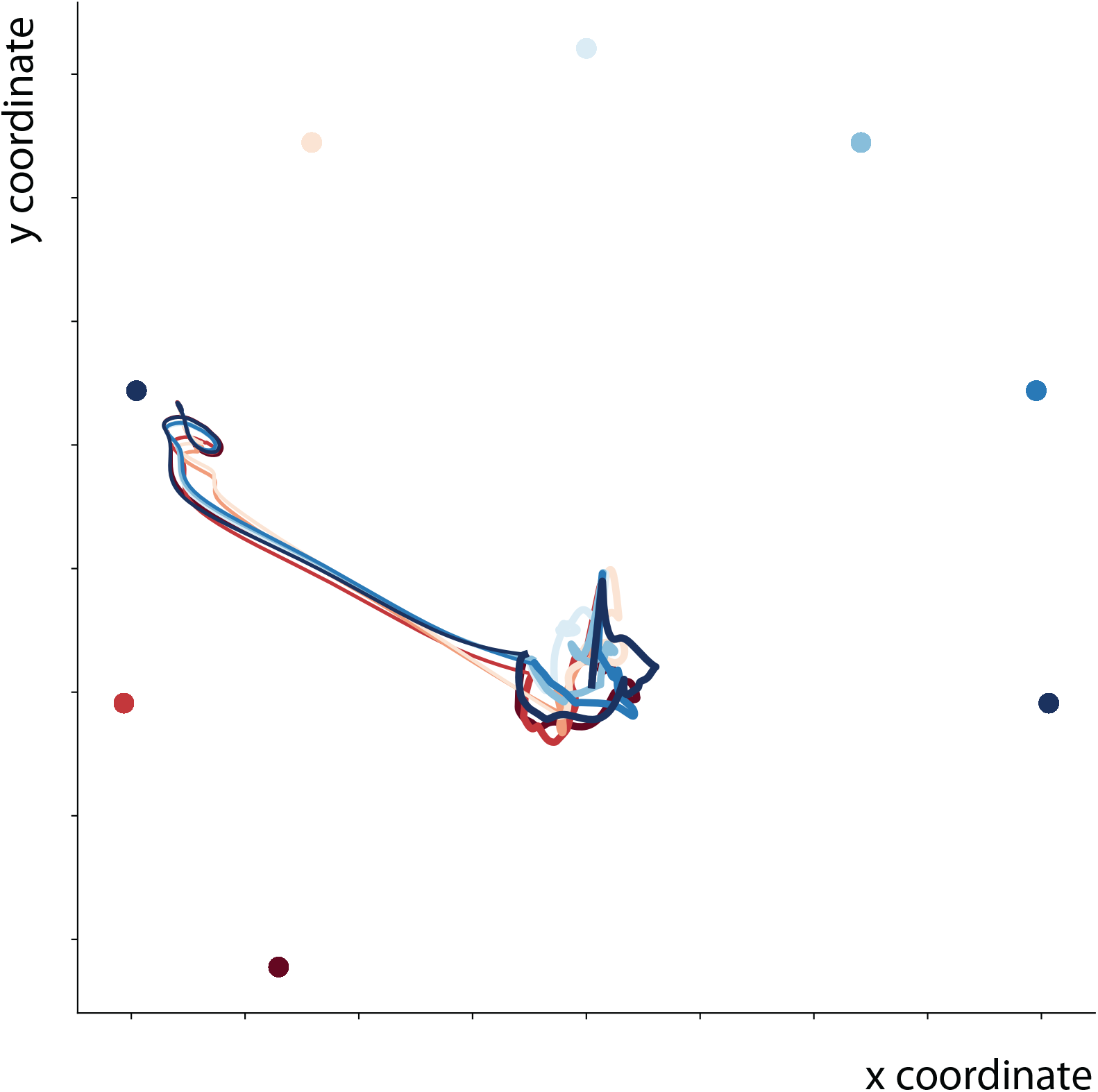
Controllers trained using DRL generalize to target switch experiments and execute accurate reaches to the final cued target. Hand trajectories (in kinematic space) during the preparatory and movement periods. Trace color indicates the initial cued target.

## Notes

### Competing Interest Statement

The authors have declared no competing interest.

### Funding

- NIH

## References

[1] Mark M Churchland et al. “Neural population dynamics during reaching”. In: Nature 487.7405 (2012), pp. 51–56.

[2] Krishna V Shenoy, Maneesh Sahani, and Mark M Churchland. “Cortical control of arm movements: a dynamical systems perspective”. In: Annual review of neuroscience 36.1 (2013), pp. 337–359.

[3] Patrick T Sadtler et al. “Neural constraints on learning”. In: Nature 512.7515 (2014), pp. 423–426.

[4] Evan D Remington et al. “Flexible sensorimotor computations through rapid reconfiguration of cortical dynamics”. In: Neuron 98.5 (2018), pp. 1005–1019.

[5] Emily R Oby et al. “Dynamical constraints on neural population activity”. In: Nature Neuroscience 28.2 (2025), pp. 383–393.

[6] Steven P Wise, Michael Weinrich, and Karl-Heinz Mauritz. “Movement-related activity in the premotor cortex of rhesus macaques”. In: Progress in brain research 64 (1986), pp. 117–131.

[7] Daniel W Moran and Andrew B Schwartz. “Motor cortical representation of speed and direction during reaching”. In: Journal of neurophysiology 82.5 (1999), pp. 2676–2692.

[8] Mark M Churchland and Krishna V Shenoy. “Delay of movement caused by disruption of cortical preparatory activity”. In: Journal of neurophysiology 97.1 (2007), pp. 348–359.

[9] Mark M Churchland et al. “Cortical preparatory activity: representation of movement or first cog in a dynamical machine?” In: Neuron 68.3 (2010), pp. 387–400.

[10] K Cora Ames, Stephen I Ryu, and Krishna V Shenoy. “Neural dynamics of reaching following incorrect or absent motor preparation”. In: Neuron 81.2 (2014), pp. 438–451.

[11] Xulu Sun et al. “Cortical preparatory activity indexes learned motor memories”. In: Nature 602.7896 (2022), pp. 274–279.

[12] Mark M Churchland et al. “Neural variability in premotor cortex provides a signature of motor preparation”. In: Journal of Neuroscience 26.14 (2006), pp. 3697–3712.

[13] Donald J Crammond and John F Kalaska. “Prior information in motor and premotor cortex: activity during the delay period and effect on pre-movement activity”. In: Journal of neurophysiology 84.2 (2000), pp. 986–1005.

[14] Silvia Arber and Rui M Costa. “Connecting neuronal circuits for movement”. In: Science 360.6396 (2018), pp. 1403–1404.

[15] Alastair J Loutit, Richard M Vickery, and Jason R Potas. “Functional organization and connectivity of the dorsal column nuclei complex reveals a sensorimotor integration and distribution hub”. In: Journal of Comparative Neurology 529.1 (2021), pp. 187–220.

[16] P Andersen et al. “Mechanisms of synaptic transmission in the cuneate nucleus”. In: Journal of Neurophysiology 27.6 (1964), pp. 1096–1116.

[17] Rafael V Bretas et al. “Secondary somatosensory cortex of primates: beyond body maps, toward conscious self-in-the-world maps”. In: Experimental Brain Research 238.2 (2020), pp. 259–272.

[18] Leah A Krubitzer and Jon H Kaas. “The somatosensory thalamus of monkeys: cortical connections and a redefinition of nuclei in marmosets”. In: Journal of Comparative Neurology 319.1 (1992), pp. 123–140.

[19] MJ Prud’Homme and John F Kalaska. “Proprioceptive activity in primate primary somatosensory cortex during active arm reaching movements”. In: Journal of neurophysiology 72.5 (1994), pp. 2280–2301.

[20] Christoph Fromm and Edward V Evarts. “Pyramidal tract neurons in somatosensory cortex: central and peripheral inputs during voluntary movement”. In: Brain research 238.1 (1982), pp. 186–191.

[21] Britton A Sauerbrei et al. “Cortical pattern generation during dexterous movement is inputdriven”. In: Nature 577.7790 (2020), pp. 386–391.

[22] Timothy P Lillicrap and Stephen H Scott. “Preference distributions of primary motor cortex neurons reflect control solutions optimized for limb biomechanics”. In: Neuron 77.1 (2013), pp. 168–179.

[23] Masaya Hirashima and Daichi Nozaki. “Learning with slight forgetting optimizes sensorimotor transformation in redundant motor systems”. In: PLoS Computational Biology 8.6 (2012), e1002590.

[24] Yuki Ueyama. “Mini-max feedback control as a computational theory of sensorimotor control in the presence of structural uncertainty”. In: Frontiers in computational neuroscience 8 (2014), p. 119.

[25] David Sussillo and Larry F Abbott. “Generating coherent patterns of activity from chaotic neural networks”. In: Neuron 63.4 (2009), pp. 544–557.

[26] David Sussillo. “Neural circuits as computational dynamical systems”. In: Current opinion in neurobiology 25 (2014), pp. 156–163.

[27] CK Chow and DH Jacobson. “Studies of human locomotion via optimal programming”. In: Mathematical Biosciences 10.3-4 (1971), pp. 239–306.

[28] Herbert Hatze and Johan D Buys. “Energy-optimal controls in the mammalian neuromuscular system”. In: Biological cybernetics 27.1 (1977), pp. 9–20.

[29] Frank C Anderson and Marcus G Pandy. “Dynamic optimization of human walking”. In: J. Biomech. Eng. 123.5 (2001), pp. 381–390.

[30] John Rasmussen, Michael Damsgaard, and Michael Voigt. “Muscle recruitment by the min/max criterion—a comparative numerical study”. In: Journal of biomechanics 34.3 (2001), pp. 409–415.

[31] Winston L Nelson. “Physical principles for economies of skilled movements”. In: Biological cybernetics 46.2 (1983), pp. 135–147.

[32] Emanuel Todorov and Michael I Jordan. “Smoothness maximization along a predefined path accurately predicts the speed profiles of complex arm movements”. In: Journal of Neurophysiology 80.2 (1998), pp. 696–714.

[33] Christopher M Harris and Daniel M Wolpert. “Signal-dependent noise determines motor planning”. In: Nature 394.6695 (1998), pp. 780–784.

[34] Marcus G Pandy et al. “An optimal control model for maximum-height human jumping”. In: Journal of biomechanics 23.12 (1990), pp. 1185–1198.

[35] Gerald E Loeb, WS Levine, and Jiping He. “Understanding sensorimotor feedback through optimal control”. In: Cold Spring Harbor symposia on quantitative biology. Vol. 55. Cold Spring Harbor Laboratory Press. 1990, pp. 791–803.

[36] Emanuel Todorov and Michael I Jordan. “Optimal feedback control as a theory of motor coordination”. In: Nature neuroscience 5.11 (2002), pp. 1226–1235.

[37] Tamar Flash and Neville Hogan. “The coordination of arm movements: an experimentally confirmed mathematical model”. In: Journal of neuroscience 5.7 (1985), pp. 1688–1703.

[38] Yoji Uno, Mitsuo Kawato, and Rika Suzuki. “Formation and control of optimal trajectory in human multijoint arm movement”. In: Biological cybernetics 61.2 (1989), pp. 89–101.

[39] Dan Liu and Emanuel Todorov. “Evidence for the flexible sensorimotor strategies predicted by optimal feedback control”. In: Journal of Neuroscience 27.35 (2007), pp. 9354–9368.

[40] Masahiko Haruno and Daniel M Wolpert. “Optimal control of redundant muscles in step-tracking wrist movements”. In: Journal of Neurophysiology 94.6 (2005), pp. 4244–4255.

[41] Emanuel Todorov and Weiwei Li. “A generalized iterative LQG method for locally-optimal feedback control of constrained nonlinear stochastic systems”. In: Proceedings of the 2005, American Control Conference, 2005. IEEE. 2005, pp. 300–306.

[42] Yuval Tassa, Nicolas Mansard, and Emo Todorov. “Control-limited differential dynamic programming”. In: 2014 IEEE International Conference on Robotics and Automation (ICRA). IEEE. 2014, pp. 1168–1175.

[43] Weiwei Li and Emanuel Todorov. “Iterative linear quadratic regulator design for nonlinear biological movement systems”. In: First International Conference on Informatics in Control, Automation and Robotics. Vol. 2. SciTePress. 2004, pp. 222–229.

[44] Olivier Codol et al. “MotorNet, a Python toolbox for controlling differentiable biomechanical effectors with artificial neural networks”. In: Elife 12 (2024), RP88591.

[45] David Sussillo et al. “A neural network that finds a naturalistic solution for the production of muscle activity”. In: Nature neuroscience 18.7 (2015), pp. 1025–1033.

[46] Ludovica Bachschmid-Romano, Nicholas G Hatsopoulos, and Nicolas Brunel. “Interplay between external inputs and recurrent dynamics during movement preparation and execution in a network model of motor cortex”. In: Elife 12 (2023), e77690.

[47] David Raposo, Matthew T Kaufman, and Anne K Churchland. “A category-free neural population supports evolving demands during decision-making”. In: Nature neuroscience 17.12 (2014), pp. 1784–1792.

[48] Kevin L Briggman and WB Kristan Jr. “Multifunctional pattern-generating circuits”. In: Annu. Rev. Neurosci. 31.1 (2008), pp. 271–294.

[49] Isaac Kurtzer, Troy M Herter, and Stephen H Scott. “Random change in cortical load representation suggests distinct control of posture and movement”. In: Nature neuroscience 8.4 (2005), pp. 498–504.

[50] Ethan M Meyers et al. “Dynamic population coding of category information in inferior temporal and prefrontal cortex”. In: Journal of neurophysiology 100.3 (2008), pp. 1407–1419.

[51] Mattia Rigotti et al. “The importance of mixed selectivity in complex cognitive tasks”. In: Nature 497.7451 (2013), pp. 585–590.

[52] Gamaleldin F Elsayed et al. “Reorganization between preparatory and movement population responses in motor cortex”. In: Nature communications 7.1 (2016), p. 13239.

[53] Matthew T Kaufman et al. “Cortical activity in the null space: permitting preparation without movement”. In: Nature neuroscience 17.3 (2014), pp. 440–448.

[54] Mark M Churchland and Krishna V Shenoy. “Preparatory activity and the expansive null-space”. In: Nature Reviews Neuroscience 25.4 (2024), pp. 213–236.

[55] Timothy P Lillicrap et al. “Continuous control with deep reinforcement learning”. In: arXiv preprint arXiv:1509.02971 (2015).

[56] Tuomas Haarnoja et al. “Soft actor-critic: Off-policy maximum entropy deep reinforcement learning with a stochastic actor”. In: International conference on machine learning. Pmlr. 2018, pp. 1861–1870.

[57] Dimitri Bertsekas. Reinforcement learning and optimal control. Vol. 1. Athena Scientific, 2019.

[58] Muhammad Noman Almani et al. “µ Sim: A goal-driven framework for elucidating the neural control of movement through musculoskeletal modeling”. In: bioRxiv (2024), pp. 2024–02.

[59] Ernest J Cheng and Stephen H Scott. “Morphometry of Macaca mulatta forelimb. I. Shoulder and elbow muscles and segment inertial parameters”. In: Journal of Morphology 245.3 (2000), pp. 206–224.

[60] Kirsten M Graham and Stephen H Scott. “Morphometry of Macaca mulatta forelimb. III. Moment arm of shoulder and elbow muscles”. In: Journal of morphology 255.3 (2003), pp. 301–314.

[61] Martyn Goulding, Tejapratap Bollu, and Ansgar Büschges. “Sensory Feedback and the Dynamic Control of Movement”. In: Annual Review of Neuroscience 48 (2025).

[62] Tucker Tomlinson and Lee E Miller. “Toward a proprioceptive neural interface that mimics natural cortical activity”. In: Progress in Motor Control: Theories and Translations (2016), pp. 367–388.

[63] Robert L Sainburg et al. “Control of limb dynamics in normal subjects and patients without proprioception”. In: Journal of neurophysiology 73.2 (1995), pp. 820–835.

[64] Yuki Ueyama. “Costs of position, velocity, and force requirements in optimal control induce triphasic muscle activation during reaching movement”. In: Scientific Reports 11.1 (2021), p. 16815.

[65] H Lalazar et al. “Population subspaces reflect movement intention for arm and brain-machine interface control”. In: bioRxiv (2019), p. 688259.

[66] Sherwin S Chan and Daniel W Moran. “Computational model of a primate arm: from hand position to joint angles, joint torques and muscle forces”. In: Journal of neural engineering 3.4 (2006), p. 327.

[67] Abigail A Russo et al. “Motor cortex embeds muscle-like commands in an untangled population response”. In: Neuron 97.4 (2018), pp. 953–966.

